# First Plant sedaDNA assemblage from California reveals ecological stability throughout 10,000 years of human presence at Lake Elsinore

**DOI:** 10.64898/2026.06.11.731745

**Authors:** Madeline Slimp, Lisa N. Martinez, Joshua D. Kapp, Matthew E. Kirby, Glen MacDonald, Don L. Hankins, Sebastian Melrose, Matt Johnson, Beth Shapiro, Rachel S. Meyer

## Abstract

As we face the sixth mass extinction, understanding how ecosystems have persisted—or collapsed—through millennia of changing climates and human activity is critical for preventing biodiversity loss. We bolstered the past 24,000 years of plant and mammal records using targeted capture of ancient sedimentary DNA (sedaDNA) from Southern California’s Lake Elsinore, a cultural center for the Payómkawichum (Luiseño), Cahuilla, and other Peoples. Our sedaDNA approach generated a diverse dataset that included 18 plant orders not previously documented from Lake Elsinore. We paired these records with local measurements and paleo evidence of fire regimes, climate, demographic history, and ethnobotanical knowledge. We find that ecological stability persisted for 10,000 years of continuous human presence, reflecting ecosystem resilience through major climatic shifts, altered fire regimes, and varying intensities of Indigenous land use. SedaDNA revealed increased availability of food, medicinal, and utilitarian plant taxa during this period of botanical stability, shedding light on ancient fire–environment–human interactions that can inform contemporary management strategies.

## I. INTRODUCTION

Addressing the current global biodiversity crisis, also called the sixth mass extinction (Barnosky et al., 2011), requires lessons learned from the Quaternary period’s glaciation events, divergent climatic regimes, and human emergence—a geological era that reveals how ecosystems survived or succumbed to millennia of environmental and societal upheaval. The late Pleistocene to Holocene transition encompasses the last mass extinction and die-off of megafauna in North America, defined by significant glaciation events, divergent climatic regimes, and the increase of humans (Figure 1). The millennia that follow witness global proliferation of trade, culture, technology (such as food production and processing tools from fires to mortars), and knowledge that interplays with climate change to ultimately shape our present environment. One major knowledge gap is specifically how much abrupt shifts in hydroclimates (Clark et al., 2012; Kirby et al., 2013; Hudson et al., 2019) and disruptions of ecological stability (Heusser et al., 2015) affect human economic resource availability (Knell et al., 2023) when humans continuously adapt management practices.

**Figure 1:**
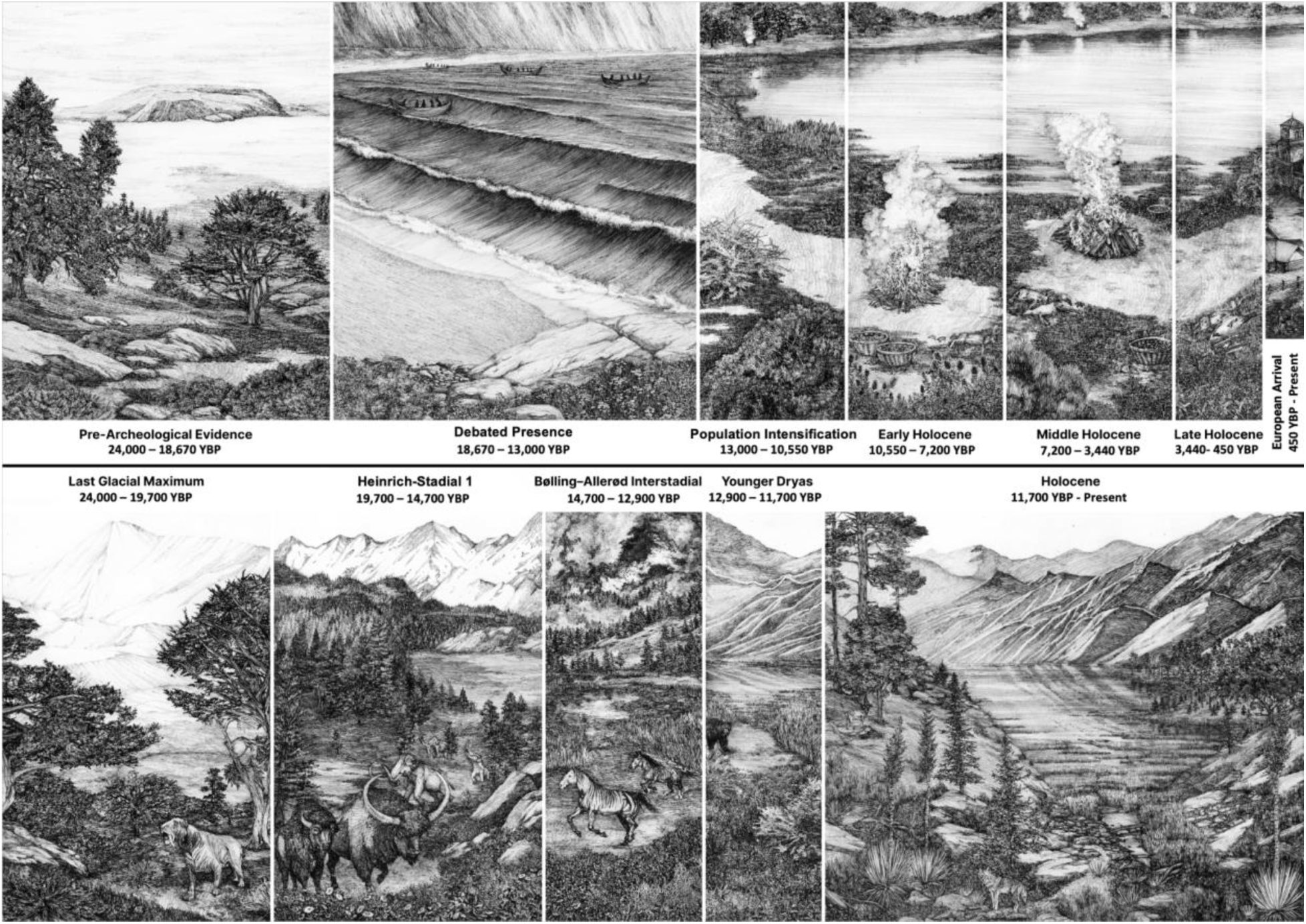
The top timeline depicts the phases of human history at Lake Elsinore used in this study. We used a combination of archaeologic, population demographic evidence, and cultural evidence in the Holocene to divide the study period into 6 subsets (See Methods: Analysis Methods). Fire depicted in this panel are small cooking fires. The bottom timeline depicts the paleoclimatic periods that occurred during the study period. Fire depicted in this panel is intense wildfire.

Sedimentary Ancient DNA (sedaDNA) can provide critical clues to describe the paleoenvironment available to humans (e.g Ekram et al., 2024 and Gelabert et al., 2025), and it is starting to be co-analyzed with traditional Indigenous knowledge and records where human practices are considered to affect the biodiversity recovered beyond faunal hunting and wildfire (Özdoğan et al., 2024; Zampirolo et al., 2024; Pears et al., 2025). Having been present before and having persisted through paleoclimatic flux (Figure 1), North American Indigenous peoples maintain oral histories that reflect these deep-time experiences rooted to their places of origin (Hankins 2024). In California, North American Indigenous peoples are credited with shaping and stewarding the landscape where, without humans, entire ecosystems would not exist (Keeley 2002; Anderson 2005; Carrico 2008; Anderson and Keeley 2018). SedaDNA has not been explored for holding valuable information about California’s past ecological communities. The warm climates of California pose a challenge because sedaDNA degrades more rapidly even when conditions are favorable, such as from lake bottoms (Lindahl and Nyberg 1974; Dabney, Meyer, and Paabo, 2013; Barnes et al., 2014). If new methods of probing highly degraded sedaDNA can reveal biodiversity change over time, such data can be interwoven with traditional knowledge to teach lessons on how climate, environment, and humans can interact. Here, we recover ancient sedimentary DNA (sedaDNA) from a well-studied Southern California Lake to fill taxonomic gaps of previous studies that recorded Southern Californian biodiversity (pollen, ethnographies, and archaeology), and to relate newfound patterns to paleoecology and to Indigenous relationships between plants and people.

SedaDNA are fragments of DNA that persist in the environment cell-free for hundreds to many thousands of years (Armbrecht et al., 2022, Kjær et al., 2022), shed by organisms through tissue loss, excrement, or decomposition. While sedaDNA is best preserved in anoxic, dry, and/or cold environments, which prevents degradation of the molecules by microbes (Dabney, Meyer, and Paabo, 2013; Barnes et al., 2014), warm climate sedaDNA, while sparser, can still offer broad taxonomic breadth than traditional methods, and is powerful in concert with other methods of paleoenvironmental reconstruction like isotope and palynology studies. In this study, we aim to use sedaDNA to generate a high confidence plant assemblage for the most recent 24,000 years of history at Lake Elsinore, a critical water resource in Southern California for millennia. The lake sits within the homeland of the Payómkawichum (Luiseño), who have stewarded its shores since time immemorial. The lake holds cultural relevance to numerous neighboring Tribes dispersed hundreds of miles from the lake itself (Figure 2). These Tribes (Juaneño, Serrano, Gabrielino, Kitanemuk, Cahuilla, and many others) did not domesticate plants in the Western sense (Meyer et al., 2012, Anderson and Wohlgemuth, 2012), but practiced cultivation and controlled burning of the land for resources (Anderson 1996 and 2006), which is hypothesized to play a key role in shaping those ancient ecologies; for example, opening up more productive grasslands than shrublands (Keeley 2002). Oral histories, archaeological investigations, and paleoenvironmental studies provide an informative patchwork of the lake’s history as an important hub for many Tribes with rich botanical resources despite a turbulent climatic history. However, they do not form a concurrent record for the Holocene when ancient human populations were the most populous and active.

**Figure 2:**
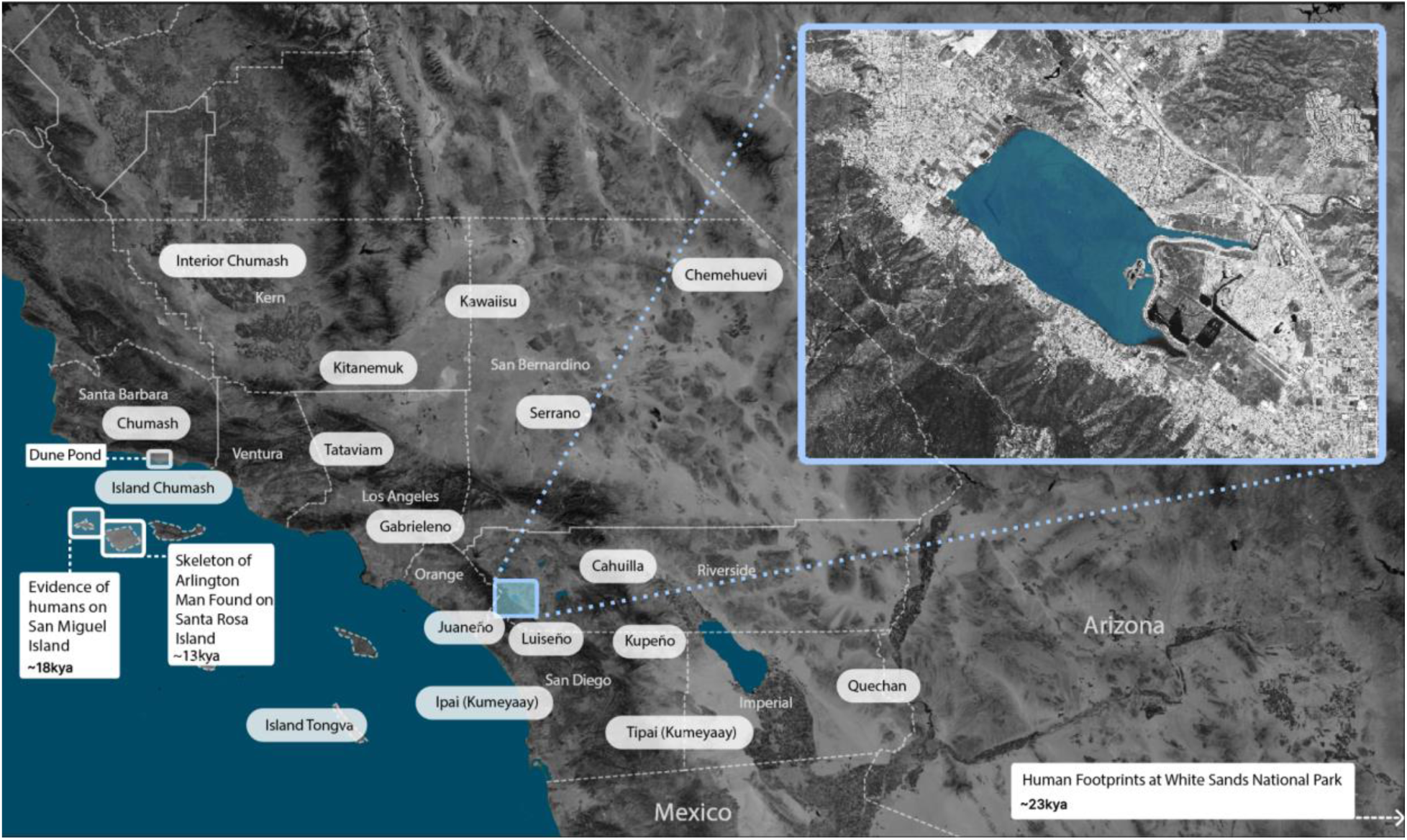
Lake Elsinore is between two well-studied and old archaeological sites to the West (Channel Islands sites) and one recently found piece of evidence of ancient humans to the East (White Sands Footprints, Bennett et al., 2021). It is located in a region once home to diverse indigenous groups who frequently interacted and traded, whose historical centers are generalized above. The box shows Lake Elsinore today, fringed by urban development and agriculture. The map of general tribal areas is based on the ‘Map of territories of Original Peoples with county boundaries in Southern California’ from the Los Angeles Almanac, 2019, which uses information from Clemmer and Heizer 1983. Dotted boundary lines are county boundaries.

We seek to connect historical plant community turnover with the historical use of plants by humans by relating sedaDNA records to human demographic estimations and ethnographies, local paleoenvironment information, and pollen records. We ask, how did communities change through time, and can these changes be linked to climatic events or human influence? What can sedaDNA add to preexisting plant assemblage reconstructions and discussions of human-plant relationships, like ethnobotanical records and archaeological remains do? Are the commercial universal target capture kits a suitable method for recovering these communities, and at a resolution fine enough to sufficiently answer these questions? Using sedaDNA from Lake Elsinore cores LEDC10-1 and LEGC03-3 spanning 24,000 years, we find that plant community shifts are largely driven by climate change and intense fire at the Pleistocene-Holocene transition, while ecological stability characterizes the most recent 10,000 years of continuous human presence. Within this stable period, human demographic phases do not drive large community-level change, but instead correspond with increases in specific economically useful taxa. We suggest that Indigenous management stabilized the local environment, and that the ecological roles and ethnobotanical practices of the Payómkawichum and other California Indigenous Peoples can be illuminated through co-analysis of sedaDNA and paleoclimate records.

## II. RESULTS

### Lake Elsinore sedaDNA recovery of broad biodiversity

Our DNA capture method using Angiosperms353 recovered a broader diversity of plant orders than traditional detection approaches. Though Angiosperms353 has been used in population-level studies (Slimp et al., 2021, Beck et al., 2021) and can be used to survey any angiosperms within reference data available for most families (One Thousand Plant Transcriptomes Initiative, 2019), this dataset was brought to the order level as a reliable taxonomic units due to the level of degradation and gaps in California-specific reference data. We recovered 50 orders in Streptophyta, the land plant phylum. Angiosperms353 target capture doubled the relative reads mapping to plants than to the shotgun approach, and had a greater taxonomic diversity (59 total orders in target capture libraries and 47 total orders in the shotgun libraries, Supp. Table 1). Even though the Angiosperms353 baits are designed to target flowering plants (Angiosperms), the probe genes were from photosynthetic pathways that persist, albeit are less conserved in mosses, ferns, and Gymnosperms. Of the 50 Streptophyta orders recovered, 13 were non-angiosperm plant orders. The 50 orders recovered with sedaDNA sum to 18 more orders than found in traditional studies, such as in pollen records, macro botanical records from archaeological sites, and ethnobotanical accounts of plants around Lake Elsinore (Figure 3). These traditional detection methods share 32 of the 50 orders captured with our sedaDNA analysis. We found that the only orders missing in our sedaDNA results that were present in pollen and ethnobotanical accounts were fern (Salvinales, Polypodiales), gymnosperm (Pinales, Cupressales, Ephedrales), or lycopod (Isoetales) orders, which were not targeted by the Angiosperms353 kit (Figure 3).

**Figure 3:**
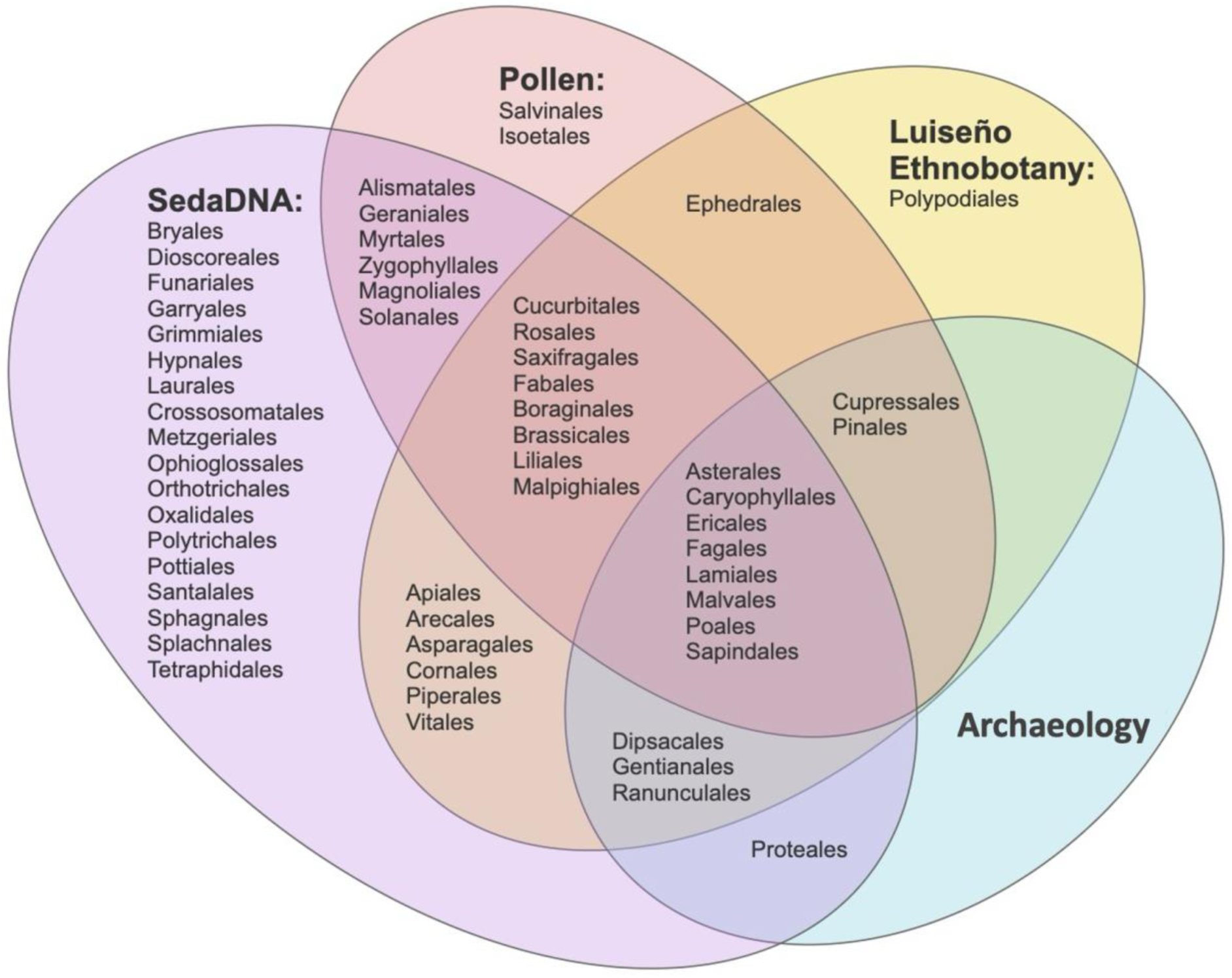
Venn Diagram that shows overlap of recorded Streptophyta orders in previous studies or literature about Luiseño ethnobotany and Lake Elsinore’s natural history. This comparison does not include a list of contemporary species in the area; it only compares quantitative and qualitative methods to reconstruct ancient plant communities. The pollen record list used pollen records published in Heusser et al. 2015, Grenada 1997, Byrne et al. 2003, and O’Keefe et al. 2023. Fossil/Archaeological records included macrofossils from the La Brea tarpits (George 2022) and floatation samples from archaeological fill (Grenda 1997). Ethnobotanical accounts for the Luiseño referenced Mead 1972, True, Meighan, and Crew 1974, Harrison 1933, Bean and Saubel 1972, Bean and Shipek 1978, and Sparkman 1908. SedaDNA data is from this study.

**Table 1:** Vegetation Shifts with Fire Regime. Pairwise Dunn tests were performed following a significant Kruskal-Wallis test. > = left category has greater abundance; < = right category has greater abundance; n.s. = not significant; “-” = Kruskal-Wallis not significant. Raw values are provided in Supplementary Document 3.

| **Plant Order** | **KW Significant?** | **Low-Fire vs. Intense Fire Transition** | **Low-Fire vs. High Fire** | **Intense Fire Transition vs. High Fire** |
| --- | --- | --- | --- | --- |
| Caryophyllales | No | - | - | - |
| Crossosomatales | Yes | n.s. | < | < |
| Alismatales | Yes | n.s. | > | n.s. |
| Fabales | Yes | n.s. | n.s. | < |
| Gentianales | Yes | > | n.s. | < |
| Rosales | Yes | n.s. | < | < |
| Brassicales | No | - | - | - |
| Myrtales | Yes | n.s. | < | < |
| Solanales | Yes | n.s. | < | < |
| Lamiales | Yes | n.s. | < | < |
| Fagales | Yes | n.s. | < | < |
| Malvales | No | - | - | - |

Green algae taxa are expected in Lake Elsinore, which experiences frequent contemporary blooms. Our target capture libraries also contained green algae (Chlorophyta), another off-target group, which fluctuated in read abundance with Streptophyta (Supplemental Figure 1). We detected 13 orders of Chlorophyta in the target capture libraries, though these were not enriched by the Angiosperms353 kit, shown by a greater proportion of Chlorophyta to total reads in the shotgun libraries (Supplemental Table 1). Chlorophyta were present in small amounts throughout the Pleistocene samples, but both Chlorophyta and Streptophyta were recovered more frequently in the Holocene. However, neither Streptophyta nor Chlorophyta total reads correlate with percent sand, the precipitation runoff proxy.

The Chlorophyta orders we recovered were a mix of expected and dubious taxonomic assignments. Of the 13 Chlorophyta orders, Eustigmatales is the most abundant order recovered, accounting for one-third of the Chlorophyta reads. Organisms in Eustigmatales are common, found at Lake Elsinore, map to expected taxa by BLASTn, and show damage patterns (Supplemental Document 1, *Nannochloropsis limnetica*). However, the second order of highest abundance is Chlorokybales, an algal clade monophyletic with land plants, which is cryptic and not expected in the region, and BLAST queries of reads that mapped to Chlorokybales match closely to many bacteria. In conclusion, much of the green algae sequence data in this study could be ancient and in high abundance, but validation is beyond the scope of this study.

Our analysis of mammal DNA using the Megamammals capture baits required careful filtering of human contamination and validation, yielding a mix of reliable ancient signals and ambiguous results. We removed any assignments to primates as they were ostensibly human sequences and found remaining reads mapped to 9 orders of mammals, with varying read depth and level of certainty about mapping accuracy because reads were short. Primate DNA subsequently removed comprised a large amount of the captured library data: 35% of the total quality-filtered reads (Supplemental Table 1; primate DNA was removed from sequence data publication to respect Tribal sovereignty over human data). In validating mammal matches, we still found some reads that mapped to non-human taxa had a top BLASTn match to *Homo sapiens* (Supplemental Table 2). Nonetheless, we did confirm some other mammal matches, such as a read matching to *Canidae* from a sample dating ∼8,900 ybp. We also verified an *Ursus arctos* (likely *Ursus arctos californicus*) read from ∼5,000 ybp and ∼6,500 ybp, and a *Felis* read from ∼5,500 ybp. Some mammal groups, like Bovidae, had both short reads that looked typical of ancient reads from expected taxa as old as ∼15,400 ybp, but also long and likely modern reads with BLAST matches to humans and other animal taxa that are common modern contaminants, like *Bos taurus* and *Sus scrofa* (Supplemental Table 2). Other patterns of DNA mapping to mammals, like bats, are interesting but contain several low to medium-complexity short reads for which we cannot rule out contamination. In summary, we did not find that the cores provided sufficient sedaDNA mammal records to look for patterns between DNA retrieval and fire, paleoclimate, or human demography.

Lake Elsinore sedaDNA is rich with sequences belonging to taxa outside of plant or mammal targets. We identified vertebrates, including birds (*Corvus*, *Meleagris*), fish (hitch-fish *Lavinia*, chub *Gila robusta*), insects, and other small fauna. Some sequences are taxonomically close to their expected species when BLAST matched. For example, sequences mapped using our closed reference database to *Gila robusta* from ∼2,260 ybp has a BLASTn match only to Cyprinidae, *Gila,* and a few close congeners. We also found that several high relative-read abundance species were detected similarly in the target capture and the shotgun datasets, such as the micro-crustacean *Daphnia,* only from the Holocene, and bacteria, such as the genus *Bacillus*, considered ubiquitous in nature, that could be focused on for other studies.

### Abundance of ancient plant reads

The 10 most abundant Streptophyta orders in our sedaDNA analysis (Poales, Caryophyllales, Crossosomatales, Asparagales, Alismatales, Fabales, Gentianales, Rosales, Brassicales, Myrtales; Figure 4) showed dynamic shifts in abundance throughout both sediment cores. DNA recovery was higher in the Holocene core, with a 3.15:1 ratio of mapped plant reads compared to the older core, likely due to better preservation in more recent sediments despite the challenging preservation conditions of warm climate lakes. Mapped reads of plant taxa were predominantly 30 bp in length across the Holocene and Pleistocene, consistent with the DNA fragmentation expected in warm-climate sedaDNA. We observed that Holocene samples with high Streptophyta read counts were often those with the best animal DNA recovery, particularly Insecta (Supplementary Figure 2).

**Figure 4:**
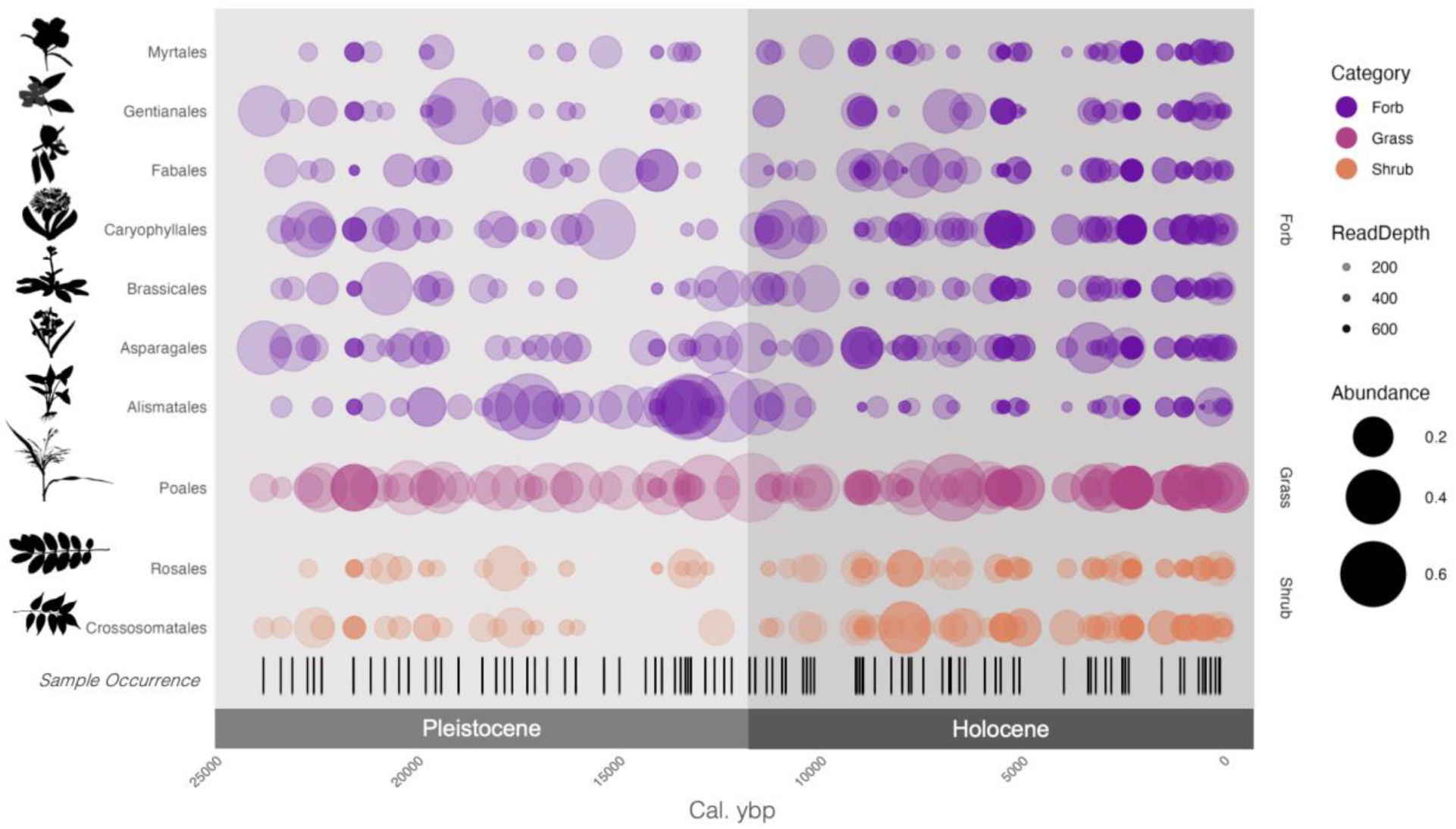
Bubble plot showing the proportion of plant reads of our samples’ top 10 most abundant plant orders. We normalized reads within samples and displayed read depth as a z-value, with the samples with the highest read depth being opaque, while the samples with smaller read depths were more transparent. The sample occurrence bar shows the sample stratification of the sediment core.

Quantification of plant sedaDNA degradation demonstrated characteristic ancient DNA damage signals throughout our lake cores. We analyzed DNA damage patterns across samples of different ages and confirmed weak to moderate damage signals characteristic of ancient DNA in the five most abundant plant orders (Supplemental Document 1). The fragment misincorporation plots show characteristic degradation patterns with elevated rates at fragment ends, though some signals appear irregular due to low read depth and sequence divergence from reference genomes. Due to the lack of a reference genome, we were unable to test Crossosomatales (the third most abundant order) and instead analyzed Fabales (the sixth most abundant order).

### Vegetation shift with fire

While long and continuous records of ancient fires are rare in coastal California, we can use published paleoproxy records to broadly categorize Pleistocene and Holocene fire regimes that pertain to our Lake Elsinore core data range from 24,000 years ago to the present. The charcoal record published in O’Keefe et al. (2023) provides a 4,000-year snapshot of fire history at Lake Elsinore in the middle of our study period (16,000-12,000 ybp; Figure 5A, Supplementary Table 3 for co-occurence with other time groupings in this study), and shows a sharp increase in charcoal accumulation rates at 13,200 ybp. We separate this elevated fire period with high charcoal accumulation and the thousand years following it into a separate period (*intense-fire transition*, 13,200-10,200 ybp) due to its distinction from the charcoal record before the elevated fire period and to look for variation in plant abundance in a window during which the ecosystem was unlikely to have reached a stable state. The *low-fire* period (24,000-13,200 ybp), which spans nearly 11,000 years of the Pleistocene, was cooler and wetter, with more infrequent fires, exemplified by low charcoal counts in sediment from nearby Baldwin Lake (Glover et al., 2020). The *high-fire* period (10,200 ybp-present) was warmer and drier with more frequent fire, with these fires fluctuating in frequency depending on vegetation type, ignition sources, and other human influence, as shown in Holocene age charcoal records from Santa Rosa Island and Dune Pond (Figure 2, Anderson et al., 2010; Ejarque et al., 2015). However, the high fire frequency in this period does not reach the amplitude of charcoal accumulation during the *intense-fire transition* period (Martinez 2020).

**Figure 5:**
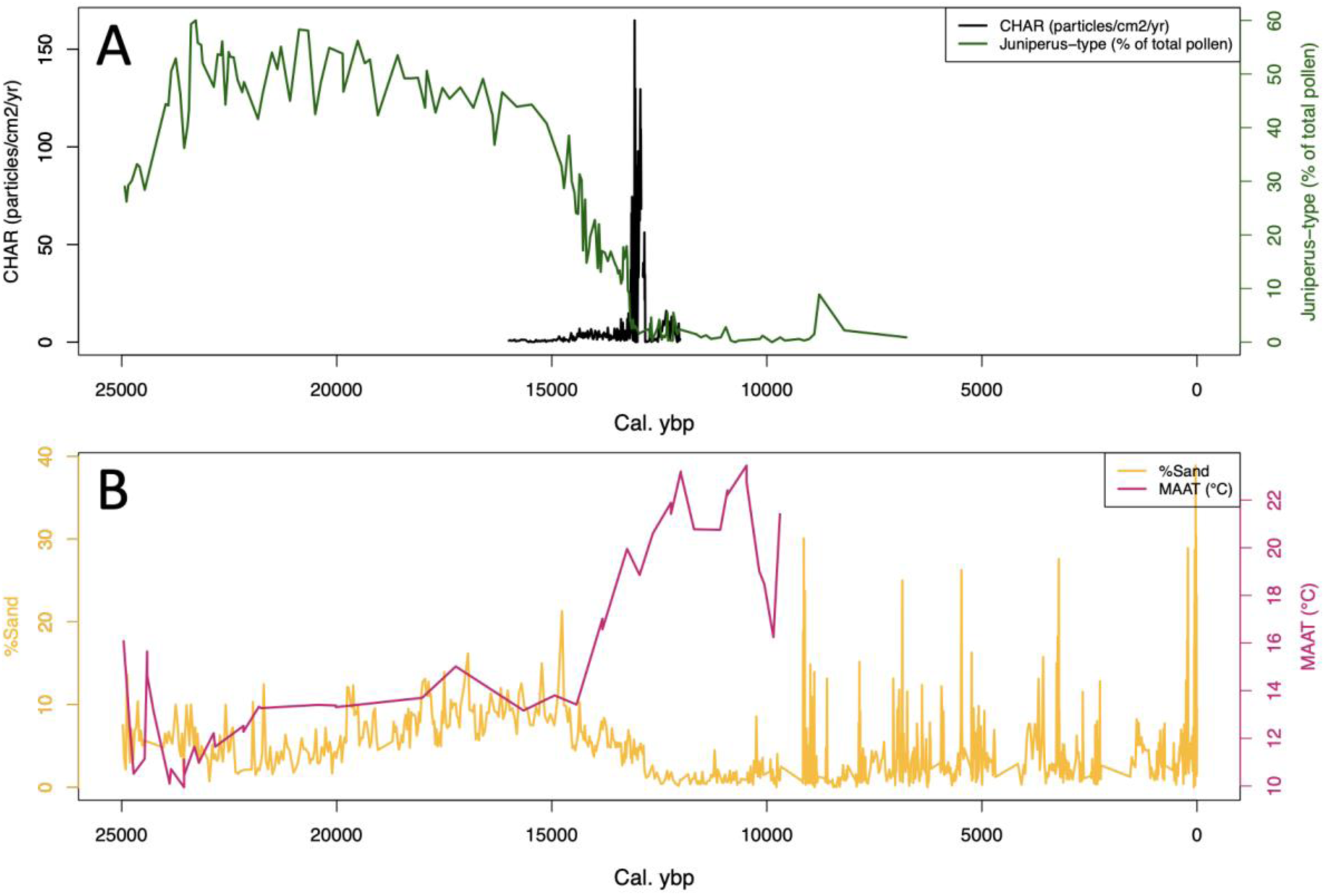
Paleoclimate and vegetation proxies from the same sediment cores collected from Lake Elsinore as used in this study. **A)** Charcoal grain counts from 16,000-12,000 cal. ybp, initially published in O’Keefe et al. (2023), is a proxy for fire. Juniperus-type pollen relative abundance, published in Huesser et al. (2015) and Kirby et al. (2018), shows gymnosperms, which were not recovered by our sedaDNA approach but are keystone in Lake Elsinore’s ancient plant communities, varied greatly in relative abundance over the study period, exemplifying high variability of the plant community. **B)** Percent sand, published in Kirby et al. (2010) using core LEGC03-3 (9,700-150 ybp) and in Kirby et al. (2018), which used core LEDC10-1 (32,000-10,000 ybp), the same cores used in this study, is a proxy for precipitation-associated runoff. This is because periods of enhanced precipitation cause greater runoff and subsequent displacement of larger sediment particles like sand in the catchment and their deposition into the lake basin. Mean Annual Air Temperature (MAAT) from 32,000-9,000 cal. ybp was published in Feakins et al. (2019).

Kruskal-Wallis and Dunn tests (Table 1) between these three fire periods and plant sedaDNA read abundance organized by plant order show that changing fire regimes significantly affect some orders but not others. The orders examined were the ten most abundant plant orders (from greatest to least read count: Poales, Caryophyllales, Crossosomatales, Asparagales, Alismatales, Fabales, Gentianales, Rosales, Brassicales, Myrtales) and four additional orders chosen because of their ethnobotanical importance in Southern California (Malvales, Solanales, Lamiales, Fagales). Orders Crossosomatales, Fagales, Myrtales, Solanales, Rosales, and Lamiales – which contain many chaparral species– maintain stable abundance from the *low-fire* to the *intense-fire transition*, then increase during the *high-fire* period (Table 1), suggesting adaptation to the high-frequency and short-interval fire regimes influenced by people. Interestingly, the grass order Poales, which also contains many chaparral taxa such as *Muhlenbergia*, *Stipa*, and *Festuca*, and riparian taxa *Typha* and *Schoenoplectus*, did not significantly fluctuate in sedaDNA over fire regime periods, nor did the Caryophyllales (order containing cacti), the Brassicales, or Malvales. The herbaceous forb order Gentianales, which includes a few chaparral species but many wetland-riparian species (Supplementary Table 4), displayed temporary decline during the *intense-fire transition* period but later returned to relative abundance levels mirroring the *low-fire* period, suggesting successful recovery (Table 1). This order did not have a significant correlation between relative abundance with percent sand, a proxy for precipitation-associated runoff and thus general winter climate wetness (Kirby et al., 2010).

The aquatic plant order Alismatales showed greater relative abundance in the Pleistocene *low-fire* period compared to the Holocene *high-fire* period (Table 1), possibly reflecting water-level variability at Lake Elsinore during the Holocene (Kirby et al., 2019, Figure 5B). However, we see no significant linear correlation between Alismatales DNA abundance and percent sand across the study period (Supplemental Figure 3). A lack of correlation may be due to two factors: the large disparity in sampling resolution between the sedaDNA and sand samples, and/or the difference in response timing and magnitude to climate between sediment deposition and vegetation.

Beta diversity analysis using Jaccard Principal Component Analysis ordination revealed a major shift in plant community composition following the *intense-fire transition* period, after which a novel Holocene assemblage persisted through the *high-fire* regime (Figure 6). Plant communities during the *low-fire* and *intense-fire transition* periods do not substantially differ in the PCoA (Figure 6A). Still, after the *intense-fire transition*, the plant community gradually shifts to a new stable state throughout the *high-fire* period (Figure 6B). This new stable state of the *high-fire* community was dominated by Poales (grasses), Fagales (tree genera including *Alnus*, *Juglans, Quercus*, and common shrubs including *Myrica*), Crossosomatales, Rosales, Myrtales (shrubs), Solanales, and Lamiales (forbs). Higher relative abundance of these orders in the *high-fire* period was supported by Dunn tests (Table 1). This community signal from sedaDNA also reflects the vegetation of the area today (Supplemental Table 4).

**Figure 6:**
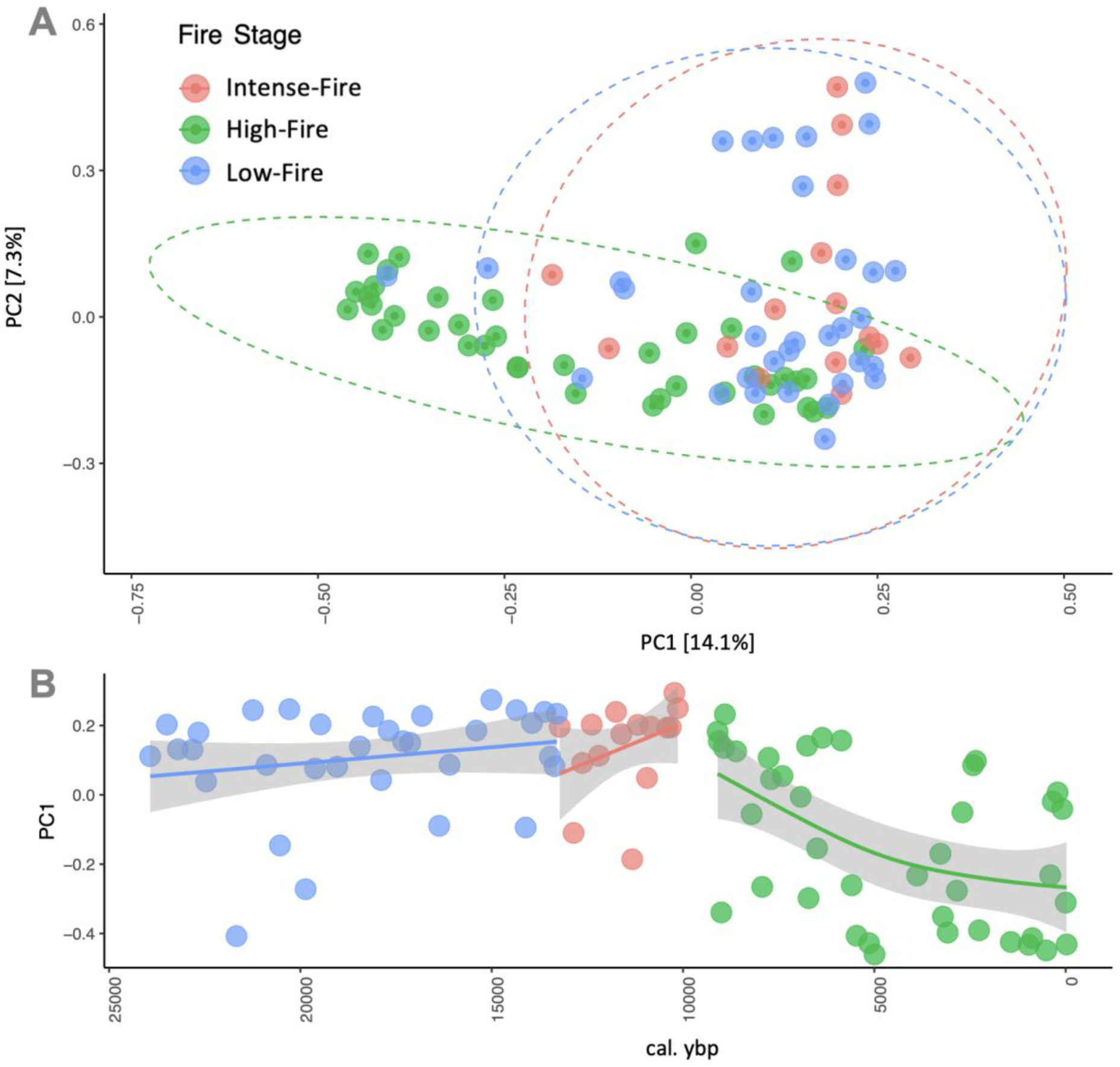
Beta diversity analysis using the Jaccard Index displayed by Fire Stage. We examined changes in the beta diversity Jaccard index, which measures community dissimilarity based on taxa presence or absence, and visualized it through ordination. **A)** PCoA space of PC1 (x-axis) vs PC2 (y-axis) **B)** PC1 (y-axis) vs time.

### Vegetation shift with paleoclimate

We found significant shifts in plant order sedaDNA relative abundance across major paleoclimate periods (Figure 1), with multiple orders shifting during the Bølling-Allerød Interstadial (14,700-12,900 ybp). The relative sedaDNA abundance of four orders significantly differed between this warm interstadial and other climatic periods occurring during the study period (Table 2). The Caryophyllales showed lower relative abundance in the Bølling-Allerød Interstadial compared to the Holocene (11,700 ybp - present) and Last Glacial Maximum (24,000-19,700 ybp). Similarly, the Crossosomatales order decreased in relative abundance during the Bølling-Allerød compared to the Holocene. In contrast, both Alismatales and Malvales exhibited greater relative abundance in the Bølling-Allerød Interstadial than in the Holocene and Last Glacial Maximum, with particularly low abundance in the Last Glacial Maximum compared to Heinrich Stadial 1 (HS-1: 19,700 - 14,700 ybp). Additionally, Alismatales maintained higher abundance in HS-1 than in the Holocene, while Malvales showed greater abundance in the Bølling-Allerød Interstadial than in the colder Younger Dryas (12,900-11,700 ybp).

**Table 2:** Vegetation Shifts with Paleoclimate. Pairwise Dunn tests were performed following a significant Kruskal-Wallis test. > = left category has greater abundance < = right category has greater abundance n.s. = not significant. Raw values are provided in Supplementary Document 3.

| **Plant Order** | **KW**<br>**Significant**<br>**?** | **Last Glacial**<br>**Maximum vs.**<br>**Bølling-Allerød**<br>**Interstadial** | **Last Glacial**<br>**Maximum vs.**<br>**Younger Dryas** | **Heinrich-**<br>**Stadial 1 vs.**<br>**Holocene** | **Bølling-Allerød**<br>**Interstadial vs.**<br>**Younger Dryas** | **Bølling-Allerød**<br>**Interstadial vs.**<br>**Holocene** | **Younger Dryas**<br>**vs.**<br>**Holocene** |
| --- | --- | --- | --- | --- | --- | --- | --- |
| Caryophyllales | Yes | > | > | n.s. | n.s. | < | < |
| Crossosomatales | Yes | > | n.s. | < | n.s. | < | n.s. |
| Alismatales | Yes | < | n.s. | > | n.s. | > | n.s. |
| Fabales | No | - | - | - | - | - | - |
| Gentianales | No | - | - | - | - | - | - |
| Rosales | Yes | n.s. | n.s. | < | n.s. | n.s. | < |
| Brassicales | No | - | - | - | - | - | - |
| Myrtales | Yes | n.s. | n.s. | n.s. | n.s. | n.s. | n.s. |
| Solanales | Yes | n.s. | n.s. | < | n.s. | n.s. | n.s. |
| Lamiales | No | - | - | - | - | - | - |
| Fagales | No | - | - | - | - | - | - |
| Malvales | Yes | < | n.s. | n.s. | > | > | n.s. |

These abundance patterns could reflect specific ecological adaptations and competitive dynamics across changing climatic conditions. Malvales, potentially represented by shrubby chaparral mallows (such as *Malacothamnus*), may have been outcompeted by other dominant chaparral taxa that became more prevalent in the Holocene, particularly Rosales and Caryophyllales, which showed increased abundance from the Younger Dryas to the Holocene (Table 2). The increased abundance of Solanales in the Holocene compared to the HS-1 suggests this order’s preference for warmer conditions (Deanna et al., 2023) in Southern California.

Plant orders show relationships with environmental proxies for moisture and lake productivity. Caryophyllales and Crossosomatales contain drought-tolerant species that could have taken advantage of drier conditions, increasing their relative abundance in the drier Holocene (Table 2; Figure 5B). We found that only Myrtales showed a significant positive linear relationship with percent sand (Supplemental Figure 4), suggesting increased abundance during wetter conditions. Total organic matter, measured as Loss On Ignition at 550°C (LOI; Dean 1974), serves as a proxy for lake productivity. Linear relationships between LOI and relative abundance revealed contrasting patterns between two orders. Asparagales, a functionally diverse order locally including both xerophytes and herbaceous perennials, showed greater relative abundance during periods of higher biological productivity, while Crossosomatales, represented locally by xerophyte genera, decreased in abundance during these same periods (Supplemental Figure 5).

### Vegetation shift with human activity

We used local archaeological and human demographic and cultural evidence (See Methods Section, Analysis Methods) to bin human activity at Lake Elsinore into 6 phases (Supplemental Table 3), finding that changes in plant DNA abundance coincide with the phases of human activity around Lake Elsinore. The majority of significant shifts occur between low and high population phases rather than within Holocene cultural phases (Table 3). Only two orders showed significant changes during the Holocene: Solanales increased in relative abundance from the *Early Holocene* to *Late Holocene*, while Fagales increased from the *Early Holocene* to *Middle Holocene*. Significant differences in plant order abundance between low population phases (*Debated Presence* and *Population Intensification*; Table 3) and later *Holocene* phases likely reflect broader climatic changes, as our analysis of fire regimes and paleoclimate periods suggested. We found that both the *Pre-archaeological Evidence* and *European Arrival* phases exhibited no significant differences in the sedaDNA abundance of plant orders compared to other human activity phases, likely due to the small sample sizes in those bins.

**Table 3:** Vegetation Shifts with Human Settlement Phases.Pairwise Dunn tests were performed following a significant Kruskal-Wallis test. > = left category has greater abundance; < = right category has greater abundance; n.s. = not significant; “-” = Kruskal-Wallis not significant. Only pairwise comparisons with at least one significant Dunn result are shown for visibility due to high number of comparisons. Significance threshold: α = 0.05 (Bonferroni-adjusted). Raw values are provided in Supplementary Document 3.

| Plant Order | KW<br>Signifi<br>cant? | Pre-Archaeologi<br>cal<br>Evidence<br>vs.<br>Debated<br>Presence | Pre-Archaeologi<br>cal<br>Evidence<br>vs.<br>Middle<br>Holocene | Debated<br>Presence<br>vs.<br>Population<br>Intensificati<br>on | Debated<br>Presence<br>vs.<br>Early<br>Holocene | Debated<br>Presence<br>vs.<br>Middle<br>Holocene | Debated<br>Presence<br>vs.<br>Late<br>Holocene | Population<br>Intensificati<br>on vs.<br>Middle<br>Holocene | Population<br>Intensificati<br>on vs.<br>Late<br>Holocene | Early<br>Holocene<br>vs.<br>Middle<br>Holocene | Early<br>Holocene<br>vs.<br>Late<br>Holocene |
| --- | --- | --- | --- | --- | --- | --- | --- | --- | --- | --- | --- |
| Caryophyllales | No | - | - | - | - | - | - | - | - | - | - |
| Crossosomatales | Yes | n.s. | n.s. | n.s. | < | < | < | < | < | n.s. | n.s. |
| Alismatales | Yes | n.s. | n.s. | n.s. | n.s. | n.s. | n.s. | n.s. | n.s. | n.s. | n.s. |
| Fabales | No | - | - | - | - | - | - | - | - | - | - |
| Gentianales | Yes | n.s. | n.s. | n.s. | n.s. | n.s. | n.s. | n.s. | < | n.s. | n.s. |
| Rosales | Yes | n.s. | n.s. | n.s. | n.s. | n.s. | < | n.s. | < | n.s. | n.s. |
| Brassicales | Yes | n.s. | n.s. | < | n.s. | n.s. | < | n.s. | n.s. | n.s. | n.s. |
| Myrtales | Yes | n.s. | n.s. | n.s. | n.s. | n.s. | n.s. | n.s. | n.s. | n.s. | n.s. |
| Solanales | Yes | n.s. | n.s. | n.s. | n.s. | n.s. | < | n.s. | < | n.s. | < |
| Lamiales | Yes | n.s. | n.s. | n.s. | n.s. | n.s. | n.s. | < | < | n.s. | n.s. |
| Fagales | Yes | n.s. | < | n.s. | n.s. | < | n.s. | < | n.s. | < | n.s. |
| Malvales | Yes | < | n.s. | > | n.s. | n.s. | n.s. | n.s. | n.s. | n.s. | n.s. |

## III. DISCUSSION

Human activity and post-LGM fluctuating temperatures, sea level, fire regimes, and precipitation patterns all helped shape ancient ecosystems into today’s biotas, from broad community ecology to population-level evolutionary trajectories and migration patterns (Cushing, 1965). However, most studies focus on paleoclimate or paleobiology alone without the integration of historical human demography, land management, and knowledge. Our examination of the last 24,000 years of Lake Elsinore vegetation seen through sedaDNA target capture uncovers ecosystem stability following the elevated fire regime and climatic upheaval of the Pleistocene-Holocene transition. Whereas O’Keefe et al. (2023) describe specific flora and fauna between 16,000 and 12,000 ybp, and Heusser et al. (2015) and Kirby et al. (2018) described flora even earlier into the Pleistocene (32,600-9,000 ybp) using pollen from the same core as our sedaDNA work (LEDC10-01), this study encompassed all of the Holocene and relates sedaDNA results to wildfire, paleoclimate, and human demographic and cultural phases. Previous sedaDNA studies have illuminated ancient community ecologies in cold or high altitude systems, like arctic lakes, the European Alps, the Himalayas, and Beringia (Alsos et al., 2015, Garcés-Pastor et al., 2022, Liu et al., 2024, and Wanket et al., 2026 respectively). Our study extends this work by being the first to apply sedaDNA and the Angiosperms353 target capture kit to ancient warm climate Southern California, extending the geographic scope of sedaDNA studies into underrepresented Mediterranean-type ecosystems.

### Ecological change at Lake Elsinore

Our sedaDNA results harmonize well with O’Keefe et al. (2023), who first reported a community turnover at Lake Elsinore at the end of the Pleistocene and defined the change in floral community composition as a state shift, which they track from 16,000 ybp to 12,000 ybp. Although our broader sampling interval of ∼272 years compared to O’Keefe’s ∼43 years limits our ability to capture immediate post-fire vegetation responses, our plant sedaDNA results support that the *intense-fire transition* period beginning 13,200 ybp served as a tipping point, after which the ecosystem established an alternative stable state as a dominantly chaparral community. However, the sedaDNA-based community composition patterns throughout the entire Holocene suggest the formation of the contemporary chaparral community took ∼5,500 years to make (Figure 6B), and has been stable for the last ∼4,500 years. This extended period of community reorganization likely reflects the entanglement between fire regimes and the broader environmental changes of the Pleistocene-Holocene transition. The state shift between low-fire and high-fire periods encompasses more than just fire frequency, representing fundamentally different ecological epochs characterized by contrasting temperature and moisture regimes, and an evolution of human stewardship of nature likely through the transformative use of fire (Williams et al. 2001; Anderson 2005; Kirby et al., 2018). Parsing fire from human activity and climate as co-drivers of vegetation change at this transition remains a challenge, as fire as an earth system itself is entangled in vegetative, human and climatic processes (Bowman et al., 2009). Our dataset captures the net ecological outcome of these interacting forces, and adds detail to the compositional story of the woodland-to-chaparral shift.

Although we did not get species-level resolution in captured plant sedaDNA from Lake Elsinore, we tracked fourteen angiosperm orders through sedaDNA analysis and added detail to previous findings about the shift from woodland to chaparral vegetation. O’Keefe et al. (2023) report changes in some overlapping taxa, such as *Quercus* (Fagales), Cyperaceae (Poales), and Amaranthaceae (Caryophyllales), but also through gymnosperms *Juniperus* and *Pinus*, which are two major woodland taxa. In their study, *Juniperus* declines on the landscape after the *intense-fire* period, *Pinus* is resilient, and Asteraceae (Asterales) expand while Amaranthaceae and Cyperaceae remain minor. Our sedaDNA results do not focus on gymnosperms but show Rosales, Fagales, Lamiales, Solanales, Myrtales, and Crossosomatales become more abundant in the Holocene *high-fire* period after the *intense-fire* period. Each of these orders contains species that are members of chaparral or scrub communities. For example, the Myrtales order contains *Camissoniopsis,* a prolific genus in Southern California, and *Lythrum californicum*, a common species in wetlands, chaparral, and many other plant communities (Supplemental Table 4). Other groups, such as aquatic Alismatales, decrease in relative abundance after the Pleistocene, and other orders show recovery following the intense-fire period (Table 1). Additionally, we found that the Poales and Asterales orders, despite rising in relative abundance following the heightened fire period in O’Keefe’s charcoal record, had no significant shift in relative abundance in our sedaDNA analysis. This could be because our sample stratification is much coarser than theirs, or because the grass species may have turned over, such as shifting between C3 and C4 species, and we don’t see those differences at the order level. In summary, our results illuminate a complex flora inclusive of many vegetation types that make up the greater watershed around Lake Elsinore. These could relate to other processes or disturbances in the Lake Elsinore watershed, such as a reported flood from 4,800 ybp (Kirby et al., 2021).

### Human legacy at Lake Elsinore

Lake Elsinore has been a critical water resource in Southern California for millennia, influencing both its ecological and cultural history. When European settlers arrived in 1542 CE, the Payómkawichum (also known as the Luiseño, San Luiseño) inhabited the area and called the lake Páayaxchi. The Lake’s surrounding springs have cultural importance as healing centers and are directly referenced in the Payómkawichum creation story (DuBois 1908). Today, the Payómkawichum are represented by the La Jolla, Pala, Pauma, Pechanga, Rincon, San Luis Rey, Soboba, and other tribal bands and are dispersed hundreds of miles around Lake Elsinore. Archaeological investigations have rediscovered semi-permanent settlements (Grenda 1997) that corroborate oral histories of local abundance in Holocene botanical resources (DuBois 1908, Sparkman 1908), and show population increases at the lake during regionally drier spells. Together these records gesture toward a rich history of human-plant interaction at Lake Elsinore, but lack the temporal resolution and taxonomic breadth to track Holocene ecological change alongside human activity, both of which our sedaDNA record provides.

The intersection of archaeological and paleoenvironmental data at Lake Elsinore demonstrates how middle Holocene drying influenced both settlement patterns and dietary adaptations. The subsistence culture of Lake Elsinore is defined by archaeologists as stable, with small game (rabbits) being the primary focus and consistent source of food at ancient settlements on the shores of Lake Elsinore. Floral resources used at these settlements were also stable throughout the Holocene, except for the additions of acorns to the diet of Lake Elsinore inhabitants in the Middle Holocene (Grenda 1997). During the *Middle Holocene* culture (7,200-3,440 ybp), a reduction in water availability caused the number of sedentary populations to increase at Lake Elsinore. This climate change is supported locally by stable oxygen isotope and percent sand data (from LEGCO3-3), suggesting that the mid-Holocene (∼8,600-5,400 ybp) was slightly drier with more intense summer evaporation (Kirby et al., 2019). Regional middle Holocene drying is supported by sediment grain analysis of a core from Silver Lake in the central Mojave Desert, which showed that the early Holocene was wetter than the middle Holocene, with conditions becoming drier 7,800-7,400 ybp (Kirby et al., 2015). This made the broadening of resource use necessary and spurred the inclusion of mortar and pestles in food-processing areas of the settlement to process acorns (Grenda 1997). By combining Indigenous ethnographic accounts, archaeological findings, and our sedaDNA data, we argue it is possible, though speculative, that shifts in relative plant abundance during the Holocene may reflect use, cultivation, or stewardship, especially in the case of the Fagales order, which includes oaks and other economic tree species.

In the case of Lake Elsinore, we see the peak of the oak order Fagales relative abundance occurring between 7,200-3,440 ybp (the *Middle Holocene*). The appearance of the mortar and pestle during the middle Holocene culture and the increase in the relative abundance of Fagales DNA may be a sign that acorns were used as food, processed at the nearby springs that feed the lake (Lightfoot and Parrish 2009; Pechanga cultural coordinator Paul Macarro, personal communication). Mensing’s (2005, 2006) reviews of the history of the oak woodlands in California report that oaks increase in abundance across sites in California during the Holocene, which could be due to climatic shifts, with cultivation likely increasing oak populations within the last few thousand years as acorns became a popular food source. Fagales sedaDNA in the Late Holocene window returns to pre-intensification levels, which aligns with archaeological reports showing late Holocene culture (3,440-170 ybp) at Lake Elsinore shifted from permanent settlements to more sporadic visits for resource procurement (Grenda 1997).

Within the Holocene, where we see ecological stabilization, the only order-level sedaDNA shifts with human cultural phases are found only in Fagales, discussed above, and Solanales, which are medicinal and culturally significant plant groups. Solanales increased in relative abundance from the Middle to Late Holocene, and chaparral and scrub taxa in this order include food, ceremonial, and medicinal plants of particular significance to the Payómkawichum, such as two native tobacco species of *Nicotiana quadrivalvis* and *N. attenuata*, *Datura wrightii,* and ingestible berries in the genus *Solanum*.

The Holocene stable state was a landscape rich in plants of direct cultural and subsistence value to the Payómkawichum and neighboring tribes. While settlements at Lake Elsinore before the state shift from woodland to chaparral have either not been discovered or do not exist, we find that many foods and medicinal plants recorded in ethnobotanical literature and macro botanical analyses of Payómkawichum archaeological sites are members of orders relatively more abundant in the Holocene. The Rosales order contains many ethnobotanically relevant taxa, such as Holly leaf cherry (*Prunus*), California blackberry and thimbleberry (*Rubus*), serviceberry (*Amelanchier*), and elderberry (*Sambucus*), for which the fruits are used for food and dyes (Sparkman 1908). Rosales also contains stinging nettle (*Urtica*), used for its fibers to create cordage, fabric, or nets for snaring fish and small game (Sparkman 1908), as well as food and medicinal properties. The Caryophyllales order also appears frequently in ethnobotanical accounts and in pollen records (Heusser et al., 2015: *Eriogonum*, *Rumex*, and *Amaranthaceae*), including foods like the prickly pear cactus (*Opuntia*), chenopods, and miner’s lettuce (*Claytonia*), and medicines like dock (*Rumex*) (Sparkman, 1908). Poales, which, while not relatively more abundant in the Holocene (Figure 4), was the most abundant plant order overall. This taxon includes grasses used for arrow shaft making (*Elymus condensatus*) and basketry (*Muhlenbergia rigens*). We detected other orders that contain taxa known to be of particular ethnobotanical importance to the Payómkawichum, like willows in the Malphigiales order, which were used in basketweaving, medicine, and building houses, and yerba mansa (*Anemopsis californica*), a powerful medicinal herb in the Piperales order, but neither of these orders was in high abundance. The convergence of sedaDNA abundance patterns across and ethnobotanical records within these orders suggests that the Holocene landscape at Lake Elsinore was shaped in part by Indigenous stewardship for the purpose of botanical richness.

### SedaDNA for constructing ancient assemblages in warm climate sediment cores

While our analysis revealed numerous new trends and contributes to the record of ancient Southern Californian plants, the use of sedaDNA is not without its current limitations. In warm climates, DNA breaks down more rapidly and is poorly preserved (Barnes et al., 2014). This fact is evident by the sparseness of plant and mammal data, lower read counts, and short sequence lengths compared to cold climate sedaDNA studies, and plant sedaDNA is not always recovered as well as pollen records for certain taxa (e.g., oaks; Hudson et al., 2022). Additionally, a recognized limitation across both contemporary and ancient eDNA studies is incomplete species representation in reference databases (Ruppert et al., 2019). In our case, 32 Californian plant families lacked available reference data at the time of analysis, limiting our taxonomic resolution, though as databases are enriched with additional Mediterranean reference sequences, this dataset can be reanalyzed. Despite these preservation challenges, warm-climate sedaDNA can still recover meaningful ecological signals, particularly when interpreted alongside established paleoenvironmental records. In the case of the Lake Elsinore sediment cores, previous descriptive and quantitative studies provide a critical context that enabled us to interpret and recognize trends that may have been overlooked or seem speculative with sedaDNA data alone. For example, we observed the Myrtales order fluctuate over the past 24,000 years, but the addition of inferred moisture variability published by Kirby et al. (2018, 2019) allowed us to connect their relative abundance with precipitation trends.

Warm-climate sedaDNA studies do not merely reconstruct botanical pasts. They reveal landscapes actively shaped by Indigenous peoples, whose knowledge systems inform our interpretations just as our data can help reckon with fractured oral histories. Advancements in sequencing, reference databases, and validation tools are rapidly enabling sedaDNA recovery in warm climates (e.g. Australia: Campbell et al., 2025), with these regions being where human cultivation and domestication of plants originated and where stewardship practices have been most prolonged (Vavilov and Dorofeev, 1992) 1926). Ignoring the profound entanglement between ancient people and plants risks missing the mechanisms, like management practices, culture, and migration, that structured past biodiversity (Anderson 2005). At Lake Elsinore, disconnecting botanical biodiversity from Indigenous fire use and plant use would have left Holocene-scale ecological trends unexplained, while erasing the contributions of the people whose stewardship fundamentally shaped California’s landscapes. Indigenous perspectives interwoven in warm-climate sedaDNA research are therefore necessary for complete historical reconstructions and a recognition that natural history in these regions is inextricably also the history of Indigenous peoples.

## IV. METHODS

### Sampling

1cm sediment plugs were obtained from different depths of the archival halves of cores LEDC10-1 (Kirby et al., 2013) and LEGC03-3 (Kirby et al., 2007). LEGC03-3 was collected in November 2003, while LEDC10-1 was collected in June 2010 - both have been stored at 4 °C at Cal State Fullerton since acquisition. We wore gowns, hair nets, masks, and double gloves, and used razor blades and plug extraction tools sterilized with 20% bleach, 70% ethanol, and DNA-Away before each sediment extraction. 1cm plugs of sediment were placed into sterile cryovials and stored in a -80°C freezer until DNA extraction. We recorded the depths of these sediment plugs and converted them to calibrated ages using the recently improved combined age model from LEGC03-3 and LEDC10-1 by Martinez et al. (2026).

We extracted and isolated DNA from 88 sediment samples using techniques optimized for the efficient recovery of short DNA fragments. For each DNA extraction, we subsampled 50mg of sediment to 2mL Eppendorf tubes, resuspended the subsamples in 1mL of lysis buffer as described in Rohland (Rohland et al., 2018), and incubated the suspensions overnight at 37°C. Following the overnight incubation, we isolated DNA from the sediment suspensions using the silica column-based method described in Rohland (Rohland et al., 2018), following the Buffer D workflow. We eluted DNA from the column with 50µL buffer EBT (10 mM Tris, 0.05% Tween-20) and quantified the DNA extractions using the Qubit 1x dsDNA HS Assay Kit (Invitrogen) and a Qubit 4 (Invitrogen). At this stage, we included 8 extraction blanks, which went through the process above simultaneously with the true samples.

### Library Preparation

We prepared single-stranded libraries using the method outlined in Kapp et al. (2021) with the modifications described in Nguyen et al. (2023). We used either 10µL or up to 4ng of DNA extract for input into each library preparation. We double-indexed and amplified the libraries using the indexing primers outlined in Kircher et al. (2012). We prepared 50µL reactions containing 20µL pre-amplified library, 25µL Amplitaq Gold 360 Master Mix (Applied Biosystems), 2.5µL 20µM i7 indexing primer, and 2.5µL 20µM i5 indexing primer. The libraries were amplified in a Bio-Rad T100 thermocycler using the following conditions: 95°C for 10m, followed by 15 cycles of 95°C for 30s, 60°C for 30s, and 72°C for 60s, followed by 72°C for 7m. We purified all amplified libraries using 60µL (1.2X) of a SPRI bead mixture, which was prepared and performed as described in Rohland and Reich (2012). We quantified the amplified libraries using a Qubit 4 and the Qubit 1X dsDNA HS Assay Kit. We visualized several amplified libraries on a TapeStation 2200 (Agilent) using the D1000 High Sensitivity ScreenTape Assay (Agilent).

### Enrichment and Sequencing for Plants and Mammals

We performed hybridization enrichment on all libraries using probes for all angiosperms (Angiosperms353; Johnson et al., 2019) and 72 mammal mitochondria following the MyBaits v3 protocol (Arbor Biosciences) using a 36-hour incubation at 65°C. We amplified the libraries using a 50µL reaction containing 15µL of the enriched library, 1X KAPA HiFi HotStart ReadyMix, 0.5µM forward primer, and 0.5µM reverse primer. The enriched libraries were amplified in a Bio-Rad T100 thermocycler using the following conditions: 98 °C for 2m, followed by 25 cycles of 98 °C for 20s, 60 °C for 30s, and 72 °C for 60s, followed by 72 °C for 5m. The post-amplified enriched library was purified using a 1.2X SPRI clean. All enriched libraries were sequenced at the University of California, Santa Cruz Ancient and Degraded DNA Processing Center using 150-cycle kits for the Illumina NextSeq 550 sequencing system.

### Bioinformatic Methods

We wanted to test the efficacy of the target capture baits to amplify sequences from taxa of interest; thus, we performed the described pipeline on both the shotgun and target capture libraries for later comparison. We used *FastP* (v0.23.1) (Chen et al., 2018) to remove adaptors and trim reads to a minimum length of 30bp and a minimum quality score of 30, and then merged the forward and reverse reads. We mapped the merged reads with *BWA aln* (v0.7.17) (Li, 2013) with a mismatch maximum of 3 (as 10% divergence is expected within family) to a reference database consisting of plant transcriptomes from the OneKP Project (J.H. Leebens-Mack et al., 2018; E.J. Carpenter et al., 2019) and eukaryote and prokaryote mito and nuclear genomes from the National Center for Biotechnology Information (NCBI). Reads mapping to plant references were predominantly 30 bp in length, which is consistent with degradation expected in warm-climate sedaDNA. This minimum length threshold is a conservative filter that prioritizes mapping specificity over read recovery.

This study leverages the universality of the Angiosperms353 bait set, a commercially available target capture kit, universal for flowering plants, capable of resolving plant diversity down to the population level (Slimp et al., 2021). Order-level resolution in our dataset reflects degradation characteristic of sedaDNA and gaps in California-specific reference data, rather than limitations of the bait set itself, and as reference databases expand, this dataset can be reanalyzed at finer taxonomic resolution. For read mapping, we used plant transcriptomes from the OneKP Project (One Thousand Plant Transcriptomes Initiative, 2019; J.H. Leebens-Mack et al., 2018; E.J. Carpenter et al., 2019) as our primary reference database. Although the OneKP dataset comprises over 1,400 green plants, it omitted 41 angiosperm families known in the California flora. We supplemented the OneKP database by adding data from NCBI RefSeq from 9 families, leaving 32 families from California not represented. However, none of these omitted families appeared in previous pollen records or ethnobotanical accounts (Supplemental Figure 3). We used *BBMask* (v38.00) (BBMap; Bushnell 2014) to mask this reference file using Shannon’s Entropy window of 0.9 due to a high amount of low-complexity sequences passing through the pipeline. We then used *bwa samse* to generate .sam alignment files, which we converted to .bams. We sorted and indexed these alignment files and then retained only mapped reads using *samtools* (v1.13) (Li et al., 2009).

We generated coverage using *BBMap* pileup.sh (BBMap; Bushnell 2014) and then assembled a dataframe with all samples with R (v.4.4.0) (R Core Team 2021). We removed reads to targets with a prevalence in the control blanks to total target reads greater than 0.2, singletons, and human contaminants. All bioinformatic methods can be found on GitHub (https://github.com/madelineslimp/LakeElsinoreSedaDNA/tree/main).

### Analyses Methods

To classify human activity at Lake Elsinore, we used local archaeological, demographic, and cultural evidence to create 6 sample subsets (Supplemental Table 3). We used the oldest radiocarbon date from Daisy Cave (CA-SMI-261), a charcoal sample dated to 18,670 cal BP (Erlandson et al., 1996), as the lower bound of the *Debated Presence* phase (18,670-13,000 ybp). We recognize that this date is not direct evidence of human presence at or near Lake Elsinore, and that the earliest well-supported archaeological evidence of humans in Southern California is ∼13,000 ybp. However, motivated by pre-Clovis evidence broadly across North America, including human footprints dated to ∼23,000-21,000 ybp at White Sands, New Mexico (Bennett et al., 2021), we defined this earlier phase to serve as an exploratory window to examine whether sedaDNA records any ecological conditions that may have characterized the landscape available to the earliest Californians. All samples older than 18,670 ybp were grouped into the *Pre-Archaeological Evidence* group. We then used the current presumed date of the Arlington Man in California, 13,000 ybp (Rick et al., 2005, Orr 1968, Erlandson 1994, Johnson et al., 2002), as the oldest timepoint of the *Population Intensification* period (13,000-10,550 ybp), where humans were known to have settled in California.

Grenda (1997) synthesizes previous archaeological investigations and Southern California ethnographies to broadly categorize the diversity of Indigenous cultures at Lake Elsinore, delineating the Holocene cultures into three periods: the *Early Holocene* (10,550-7,200 ybp), *Middle Holocene* (7,200-3,440 ybp), and the *Late Holocene* (3,440-168 ybp). While the first evidence of housing structures on the shores of Lake Elsinore is dated to ∼9,400 ybp, we use the delineations set by Grenda (1997) to characterize human activity in the area not to discount transitory human presence at the Lake before this specific site. Archaeological evidence of housing and technology from these three periods suggests cultural changes coinciding with the three cultural periods in Southern California. Visits for resources by other neighboring tribes (Figure 2) could have also happened before the establishment of the settlement seen at Lake Elsinore. Timing of European colonization truncates the final Holocene period, the Late Holocene, at 450 ybp, making our final subset category, *European Arrival* (450-0 ybp). These cultural delineations are not drawn from Indigenous cultural concepts.

Using the published charcoal count data from Martinez (2020) and O’Keefe (2023), we classified fire activity into three groups designed around the distinct charcoal peak ∼13,200 ybp (Figure 5A; Supplemental Table 3). We binned all samples prior to the onset of elevated charcoal levels, ∼13,250 ybp, into the *low-fire* group. Samples from the elevated fire period and 2,000 years following this major event, 13,250-10,000 ybp, were binned into the *intense-fire transition* group. All younger samples were binned into *high-fire*. We also broadly group paleoclimate periods (Figure 1; Supp. Table 2), with the local LGM ending at 19,700 ybp (Kirby et al., 2018), leading into Heinrich-Stadial 1 before the Bølling–Allerød interstadial begins at 14,700 ybp. The Bølling–Allerød interstadial is punctuated by the Younger Dryas, which occurred from 12,900 ybp to 11,700 ybp. We classify 11,700 ybp to the present as the Holocene.

We recovered 50 orders in Streptophyta after removing 12 orders that did not occur in California, using https://Calflora.org to check for native species within each group. These 12 orders had low read counts, likely attributable to reference bias or mismapping. An example is 29 reads mapped to the order Welwitschiales, which has only one species in the order with its native range in Namibia. Likely, these reads are from the order Ephedrales, with which Welwitschiales shares the division Gnetophyta. There are 7 native species of *Ephedra* in California (CalFlora), but no genetic reference data for captured loci are available at the time of analysis for any species in Ephedrales. The order Ephedrales was reported in a pollen record of Lake Elsinore as existing in the area (Heusser et al., 2015) and was also recorded in Luiseno ethnobotanical accounts (True, Meighan, and Crew, 1974). While these 29 reads are likely informative, we discarded them and all reads assigned to other non-California native orders to maintain only high-confidence taxonomic assignments for comparison against pollen and ethnobotanical records. However, we did include these 12 non-native orders with dubious assignments in other downstream analyses where taxonomic accuracy is not critical, and did let them contribute to total Streptophyta read counts per sample.

We reduced the dataset to Streptophyta targets only and then binned all rows to the Order level. We normalized reads within each sample and then sorted by total read count per order, selected the top 10 plus 5 other orders of interest, and binned all other orders into ‘OTHER.’ We added these 15 orders and ‘Others’ to a data frame with all samples placed in their corresponding time period for the human activity, fire, and paleoenvironment stage (Supplemental Table 3). We performed a Kruskal-Wallis test (R Package *stats*; R Core Team 2021), using a p-value threshold of 0.05, and then performed a post-hoc Dunn Test (R Package *dunn.test*; Dinno 2014) using the Bonferroni correction model in R (v.4.4.0) (R Core Team 2021) and assigned significance to all adjusted Dunn Test P-values of less than 0.05.

To calculate beta diversity, we used the full non-normalized Streptophyta dataset (read counts for all orders per sample) to generate phyloseq objects (R Package *phyloseq* (v1.41.1, McMurdie and Holmes 2013), *ranacapa* (v0.1.0, Kandlikar et al., 2018), and *vegan* (v2.6.8, Oksanen et al., 2023)). We made the ordination with *phyloseq* (v1.41.1, McMurdie and Holmes 2013), and plotted the eigenvalues in PCA space using *ggplot2* (v3.5.1, Wickham 2016). We then plotted PC1 on the Y axis with the X axis being sample age using *ggplot2 (*v3.5.1, Wickham 2016), and used the R package *mgcv* (v1.9.1, Wood 2011) to fit a generalized additive model to the data.

### Verification of Ancient Reads

We used fastp (v.0.23.4) (Chen et al., 2018) to trim, low complexity, and quality filter the forward and reverse reads, leaving them unmerged. We then used Bowtie2 (v.2.4.4) (Langmead and Salzberg 2012) to build reference databases for select whole genomes from NCBI. We selected representative reference genomes for plant orders with total read counts above 400. Below this threshold, samples had far too few reads per sample to test, which causes jagged and uninformative plots. While the order Crossosomatales had the third highest read count, there was no reference genome available on NCBI within that order to use. We tested five orders: Poales, Caryophyllales, Asparagales, Alismatales, and Fabales. For Poales, we selected the *Juncus effusus* genome (Genbank assembly GCA_027726005.1) as the target the majority of the Poales reads mapped to was to *Juncus inflexus*, which is non-native to Southern California, where *J. effusus* is common. For Caryophyllales, we selected *Cereus fernanbucensis* (Genbank assembly GCA_024363205.1), as the highest mapped target was to a non-native member of the *Cactaceae*, while the genus *Cereus* is found in the state. For Asparagales, the most mapped to target was in Orchidaceae, and thus we selected the *Platanthera zijinensis* reference genome (Genbank Assembly GCA_039513925.1), which, while non-native to California, is a member of the only native genus available on NCBI. For the Alismatales, *Zostera marina*, an ocean plant, was the target most mapped to. Since this is not found at Lake Elsinore, we selected a reference genome that was the closest relative to *Zostera* while also being native and found in freshwater, *Triglochin maritima* (Genbank Assembly GCA_001185155.1). Lastly, for the Fabales order, we selected *Trifolium repens* (Genbank Assembly GCA_030408175.1) as a native and common substitute for *Glycine soja*. We used Bowtie2 (v2.4.4) (Langmead and Salzberg 2012) to map the trimmed reads from a selected sample to the reference genome and then used samtools (v1.13)(Li, Heng, et al., 2009) to convert the .sam file to a .bam file and then sort and index the .bam. We used Picard Tools (v3.0) (“Picard Toolkit,” 2019) to add read groups and then used mapDamage (v. 2.3.0) (Jónsson et al., 2013) to generate fragment misincorporation plots.

## Acknowledgments

This project was supported by the National Science Foundation (GSS-1759756). We thank Dr. Lisa Woodward at the Pechanga Band of Luiseño Indians’ Cultural Resources Department and Paul Macarro at the Pechanga Tribal Government Center (Payómkawichum) for several rounds of feedback on the manuscript. We thank culture bearers of the Cahuilla/ Payómkawichum people, Connor Magee, Natural and Working Lands Specialist from the Pala Band of Mission Indians, and Will Madrigal Jr., Tribal Capacities and Partnerships Program Manager for their review of this work. We also thank Dr. Ren Larison and Dr. Thomas Smith for their guidance and review of this work.

## Author Contributions

Conceptualization: RSM, BS, GM, DH, MK

Methodology: RSM, JK, LM, GM, MK

Analysis and Visualization: MS, RSM, SM

Writing—original draft: MS, RSM

Writing—review & editing: All authors

## Data Availability Statement

Raw sequencing reads for all shotgun and targeted capture libraries are publicly available in the NCBI Sequence Read Archive under BioProject accession **PRJNA1465454**. Prior to submission, reads were adapter-trimmed and quality-filtered using FastP (minimum length 30bp, minimum quality score 30), and human-derived reads were removed by alignment to the human reference genome (GRCh38.p13) using *BWA aln* with default mismatch parameters. All bioinformatic methods can be found on GitHub (https://github.com/madelineslimp/LakeElsinoreSedaDNA/tree/main).

## Supplemental Figures and Tables

**Supplemental Table 1:** Summary mapping statistics following decontamination and mapping, not including blanks. TC = target capture, SG = shotgun library before capture. The target capture libraries had more mapped reads than shotgun libraries and more plant and mammal targets that reads were mapped to, so we used this dataset (Supplementary Document 3) for analyses. While our Target Capture approach was intended for the recovery of flowering plants, increasing the proportion of Angiosperm reads by 2.4 fold when comparing to the Shotgun libraries, we mapped to taxa outside of Angiosperms, such as mosses, an abundance of alga, ferns, and gymnosperms. The Target Capture libraries had a 1.7x increase in the proportion of Gymnosperm reads. Still, a 0.2x decrease in the proportion of fern reads, a 0.6x decrease in the proportion of moss reads, and a 0.2x decrease in the proportion of Liverwort read

|  | **Target Capture Library Pool** | **Shotgun Library Pool** |
| --- | --- | --- |
| **Total reads QC passed** | 146982143 | 160297544 |
| **Total mapped reads** | 121561 | 106558 |
| **Average mapped reads per sample** | 1381.375 | 1224.804598 |
| **Percent reads mapped** | 0.08270460446 | 0.06647512953 |
| **Mapped Read Total after filtering**<br>**(Removed prevalence >0.20 and**<br>**singletons)** | 116994 | 102358 |
| **Streptophyta Targets mapped** | 360 | 159 |
| **Total targets mapped** | 883 | 586 |
| **Mammal Targets mapped (Excluding**<br>**reads mapped to human)** | 73 | 16 |
| **Average reads per target** | 137.6681767 | 174.6723549 |
| **Total mammal reads** | 1478 | 311 |
| **Proportion mammal reads to total**<br>**mapped** | 0.01215850478 | 0.002918598322 |
| **Total Streptophyta reads** | 9321 | 4416 |
| **Total Streptophyta Orders** | 59 | 47 |
| **Proportion Streptophyta reads to**<br>**total mapped reads** | 0.07667755283 | 0.04314269525 |
| **Proportion Streptophyta targets to**<br>**total targets mapped** | 0.4077010193 | 0.271331058 |
| **Total Human Contamination Reads** | 42628 | 443 |
| **Proportion human reads to total**<br>**mapped** | 0.3506716792 | 0.004157360311 |
| **Total Green Algae Reads** | 471 | 1244 |
| **Total Green Algae Orders** | 13 | 11 |
| **Total Green Algae Targets Mapped** | 20 | 35 |
| **Proportion Green Algae reads to total filtered mapped reads** | 0.004025847479 | 0.0121534223 |
| **Total Angiosperm Reads** | 8475 | 3132 |
| **Proportion Angiosperm reads to total mapped reads** | 0.06971808392 | 0.02939244355 |
| **Total Gymnosperm Reads** | 168 | 88 |
| **Proportion Gymnosperm reads to total mapped reads** | 0.001382022195 | 0.0008258413259 |
| **Total Fern Reads** | 37 | 174 |
| **Proportion Fern reads to total mapped reads** | 0.0003043739357 | 0.001632913531 |
| **Total Moss Reads** | 619 | 934 |
| **Proportion Moss reads to total mapped reads** | 0.005092093681 | 0.008765179527 |
| **Total Liverwort Reads** | 22 | 88 |
| **Proportion Liverwort reads to total mapped reads** | 0.0001809790969 | 0.0008258413259 |

**Supplemental Table 2:**
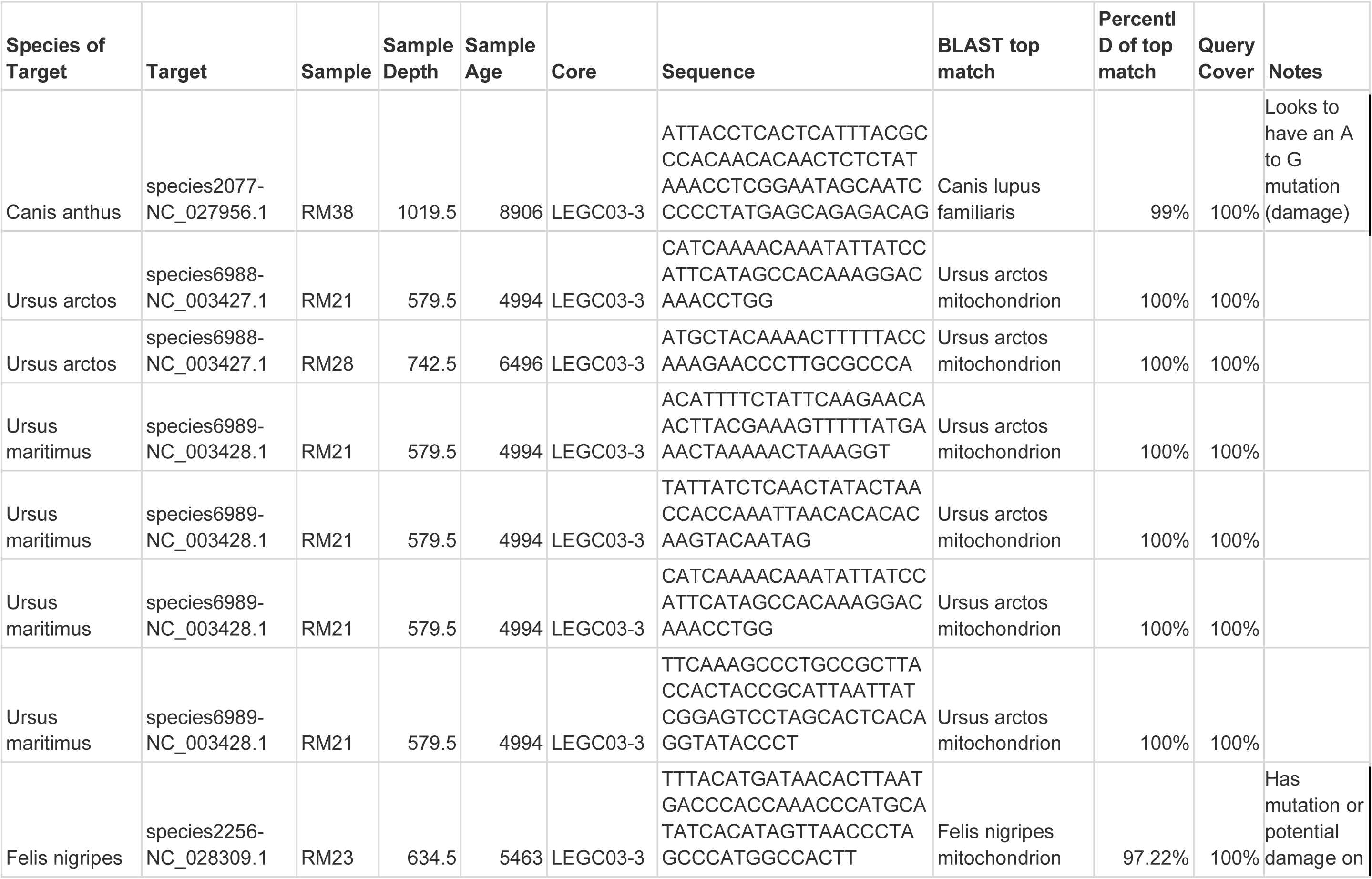
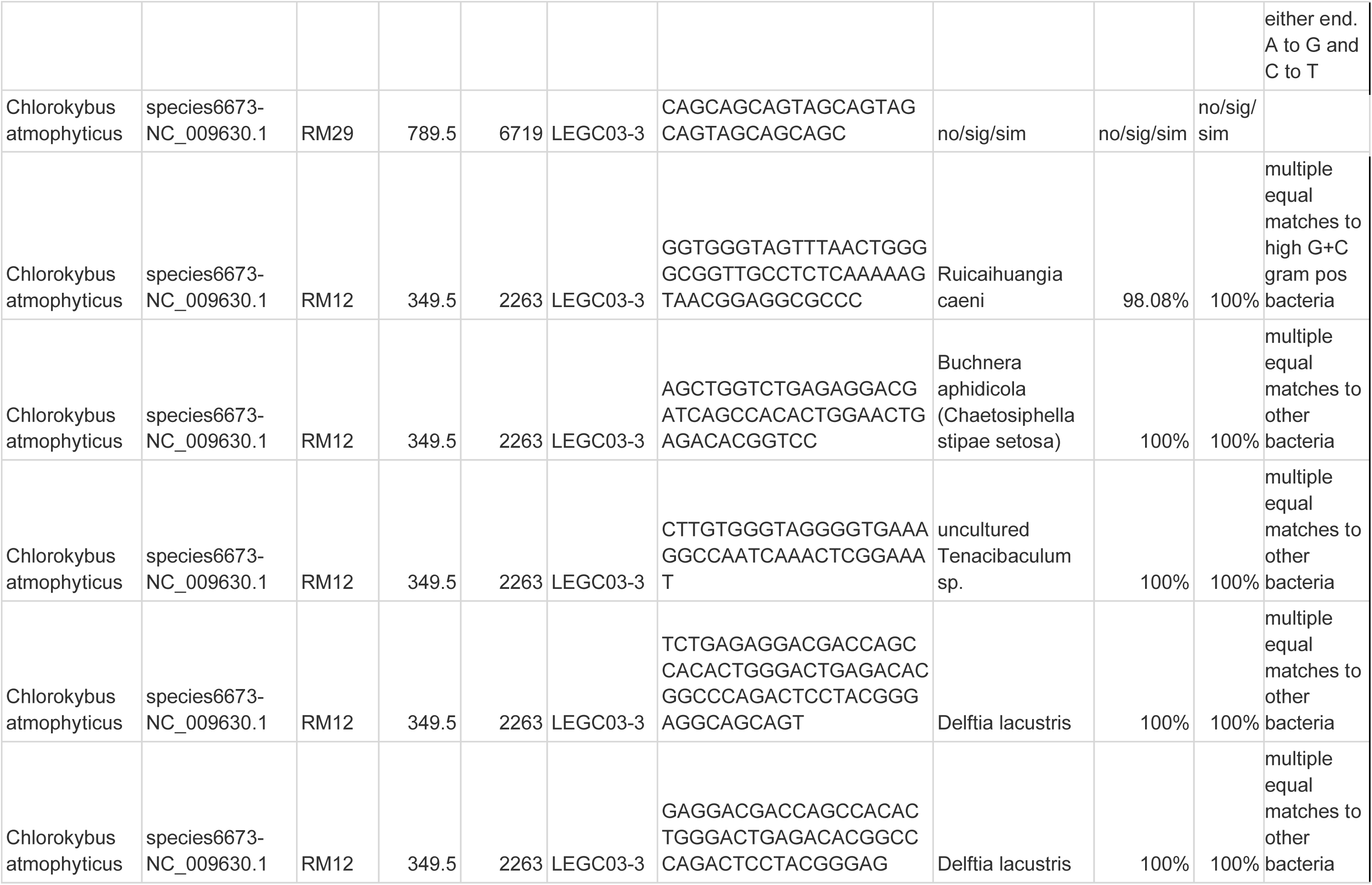
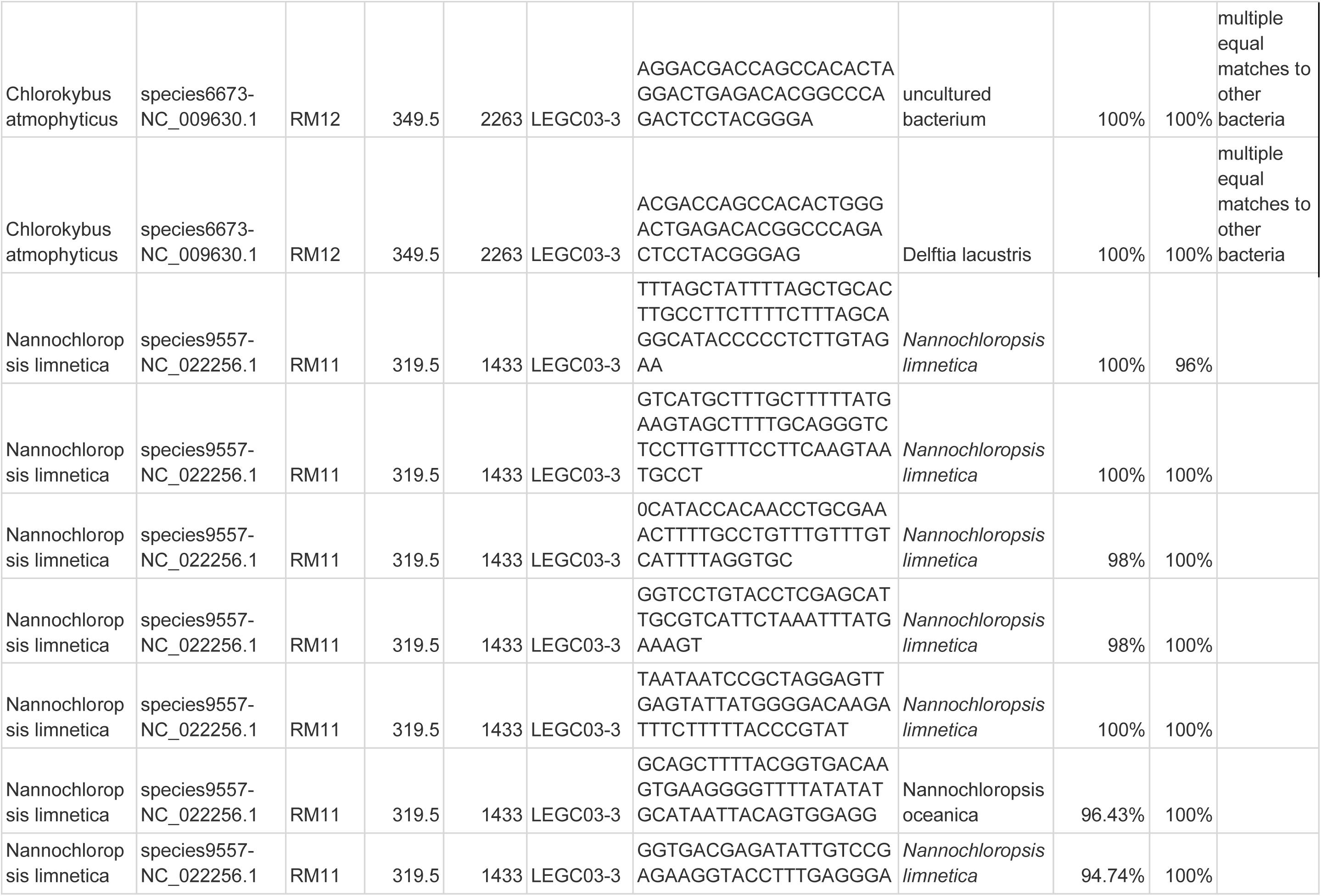
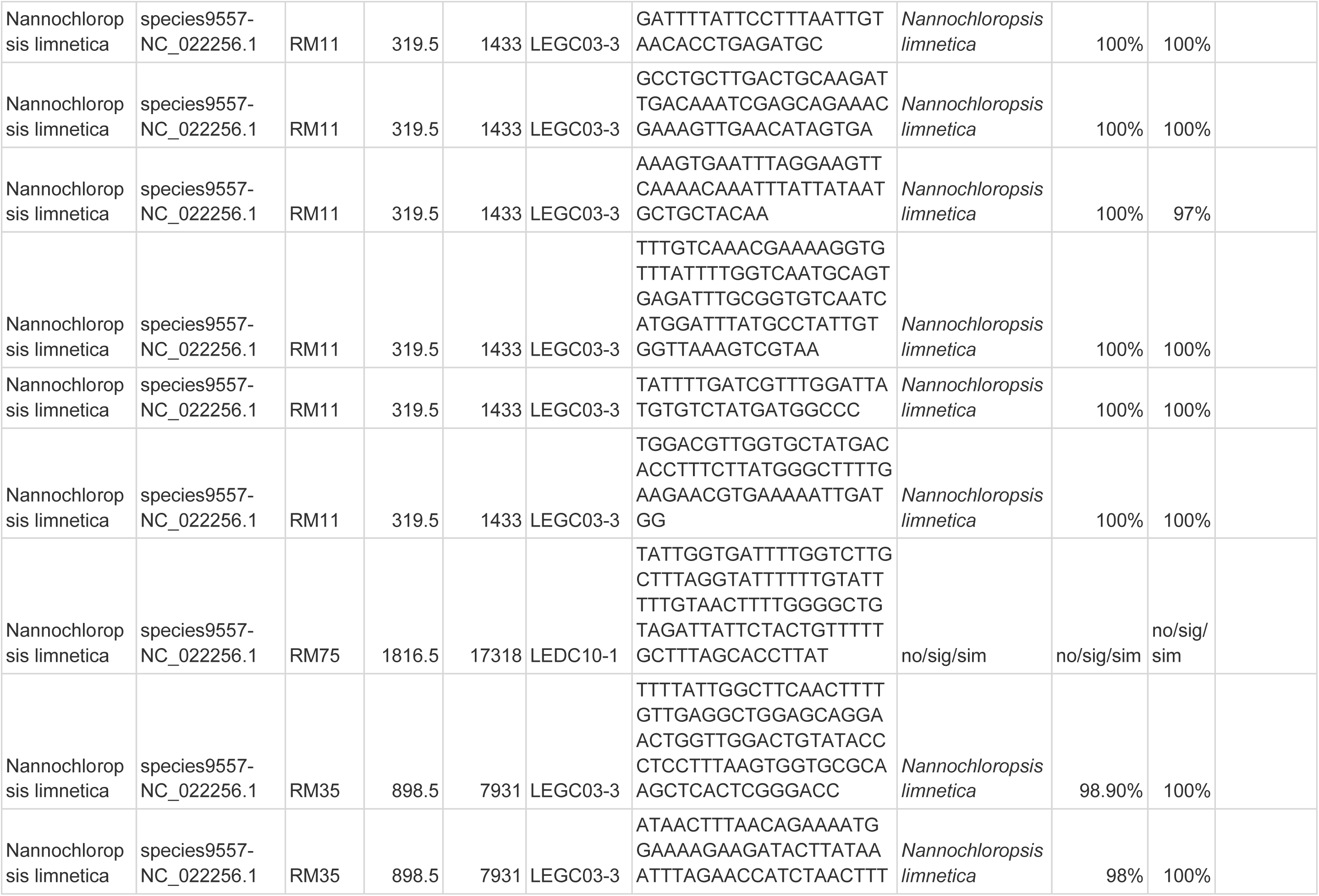
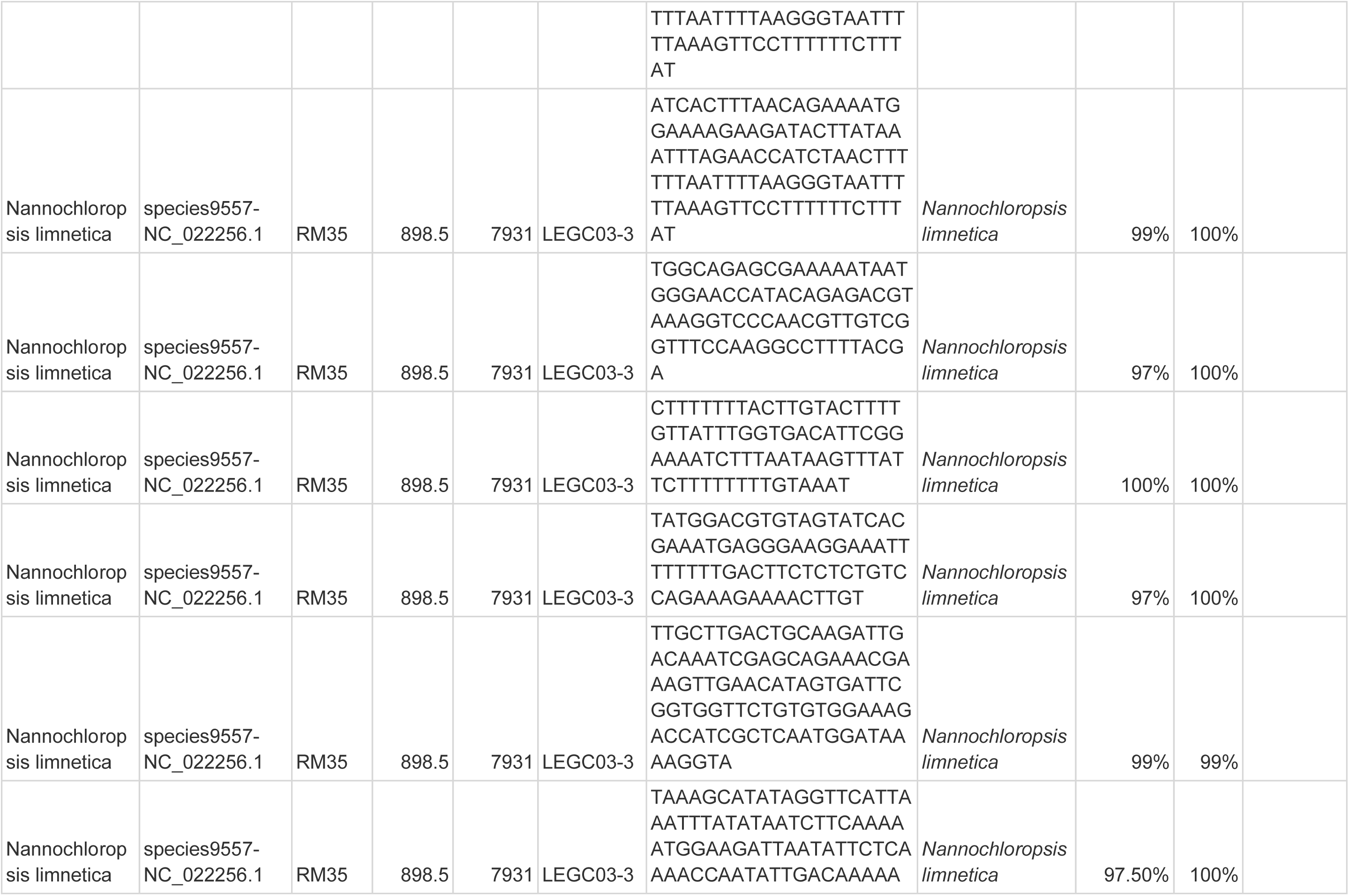
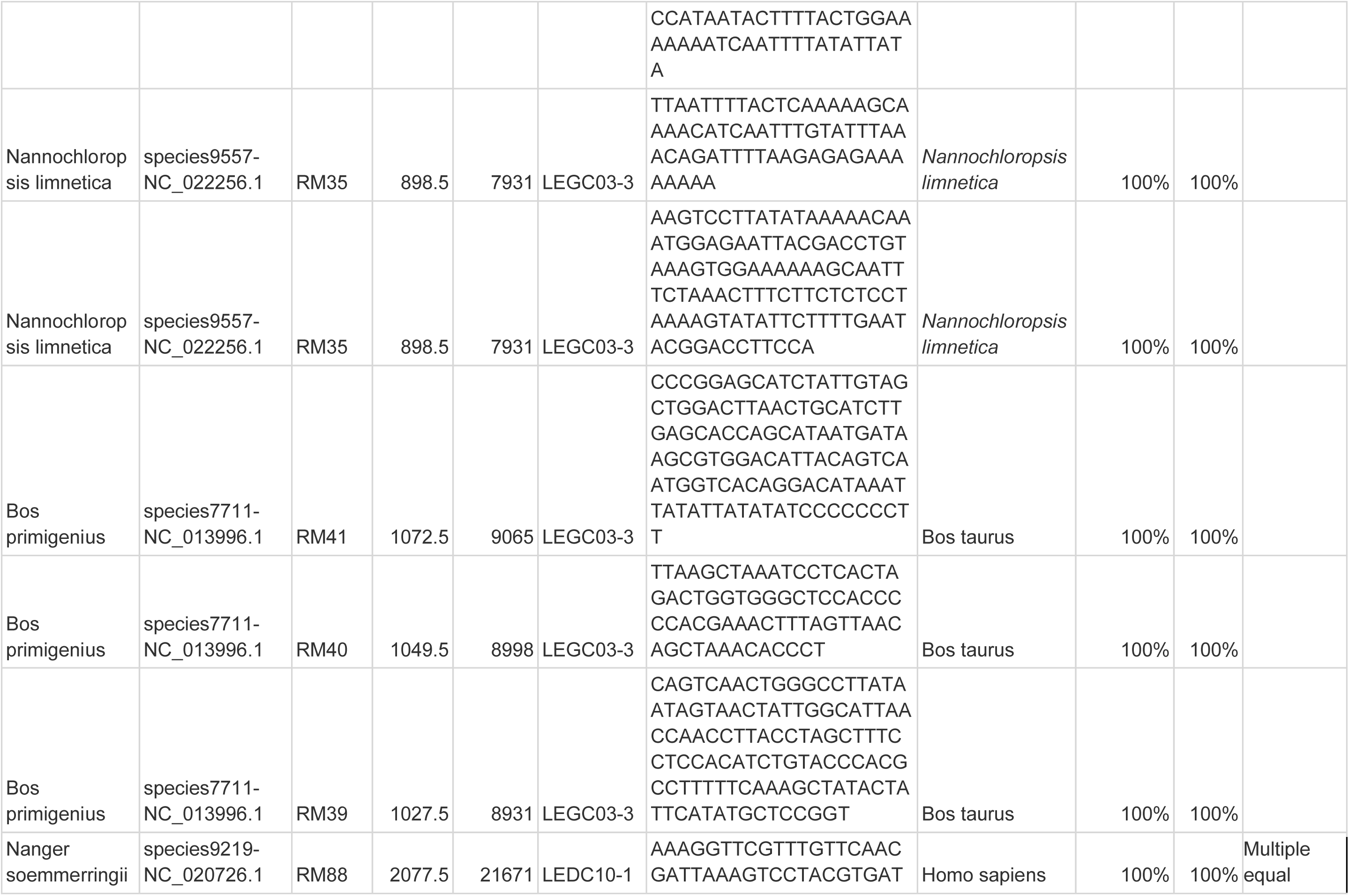
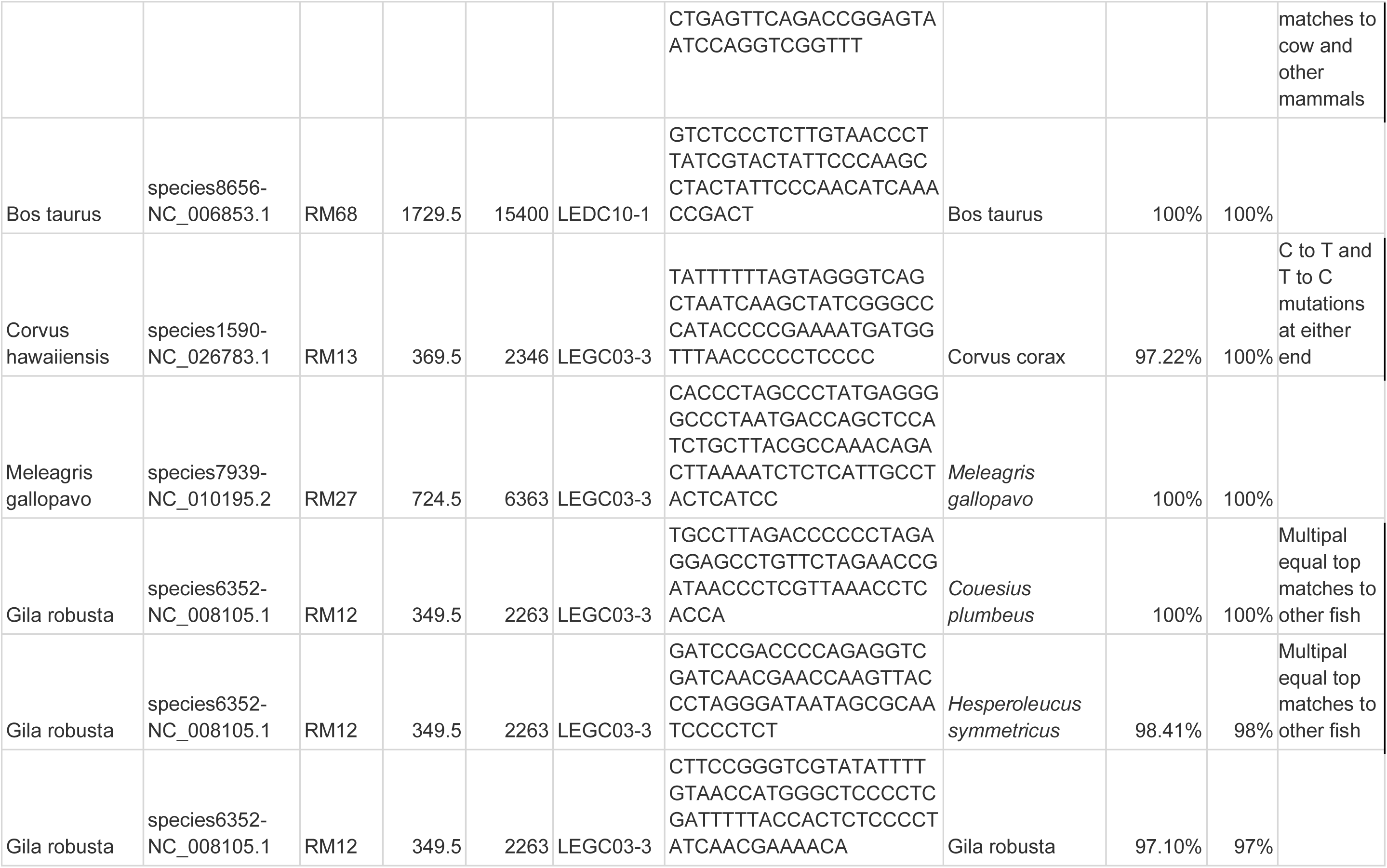
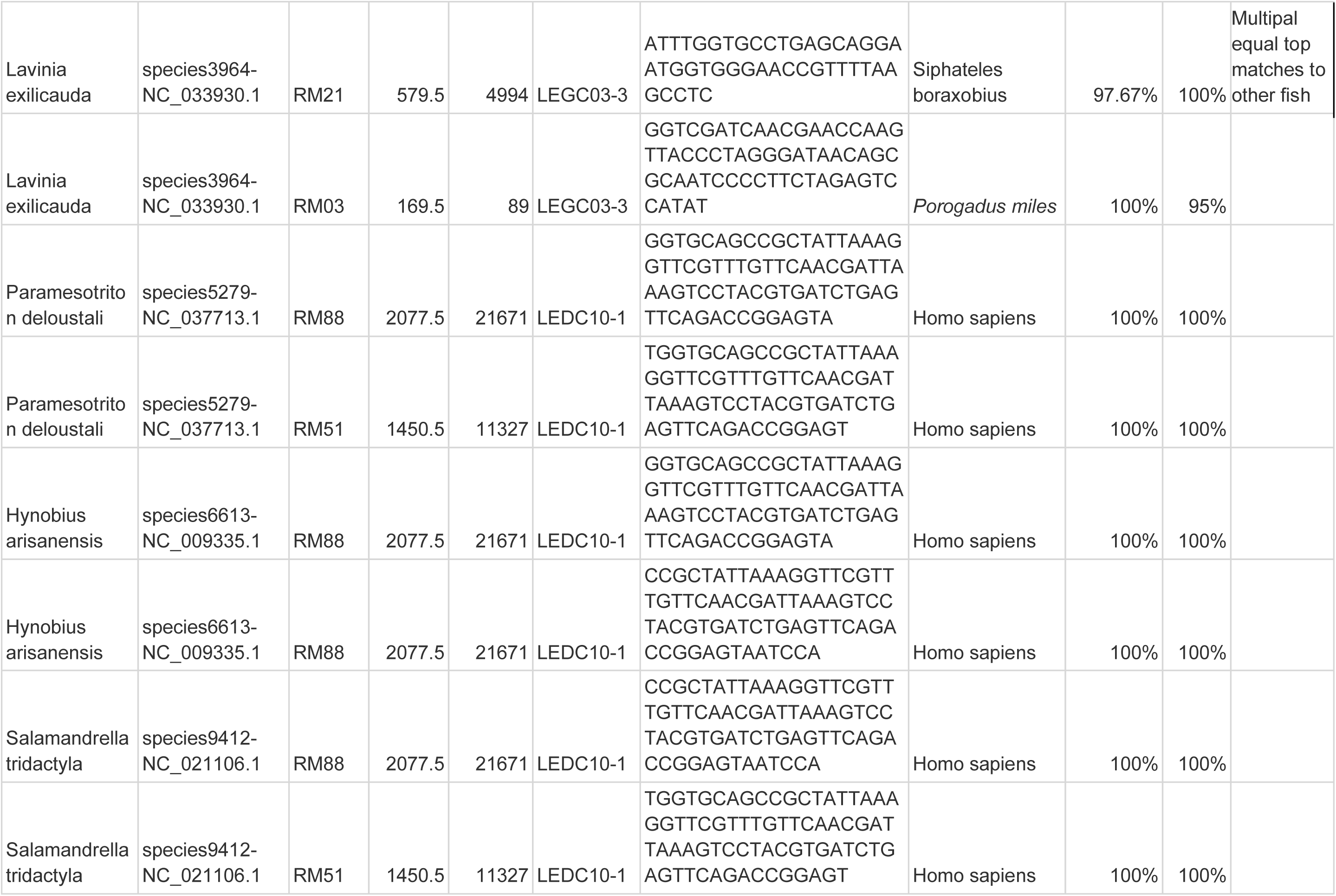
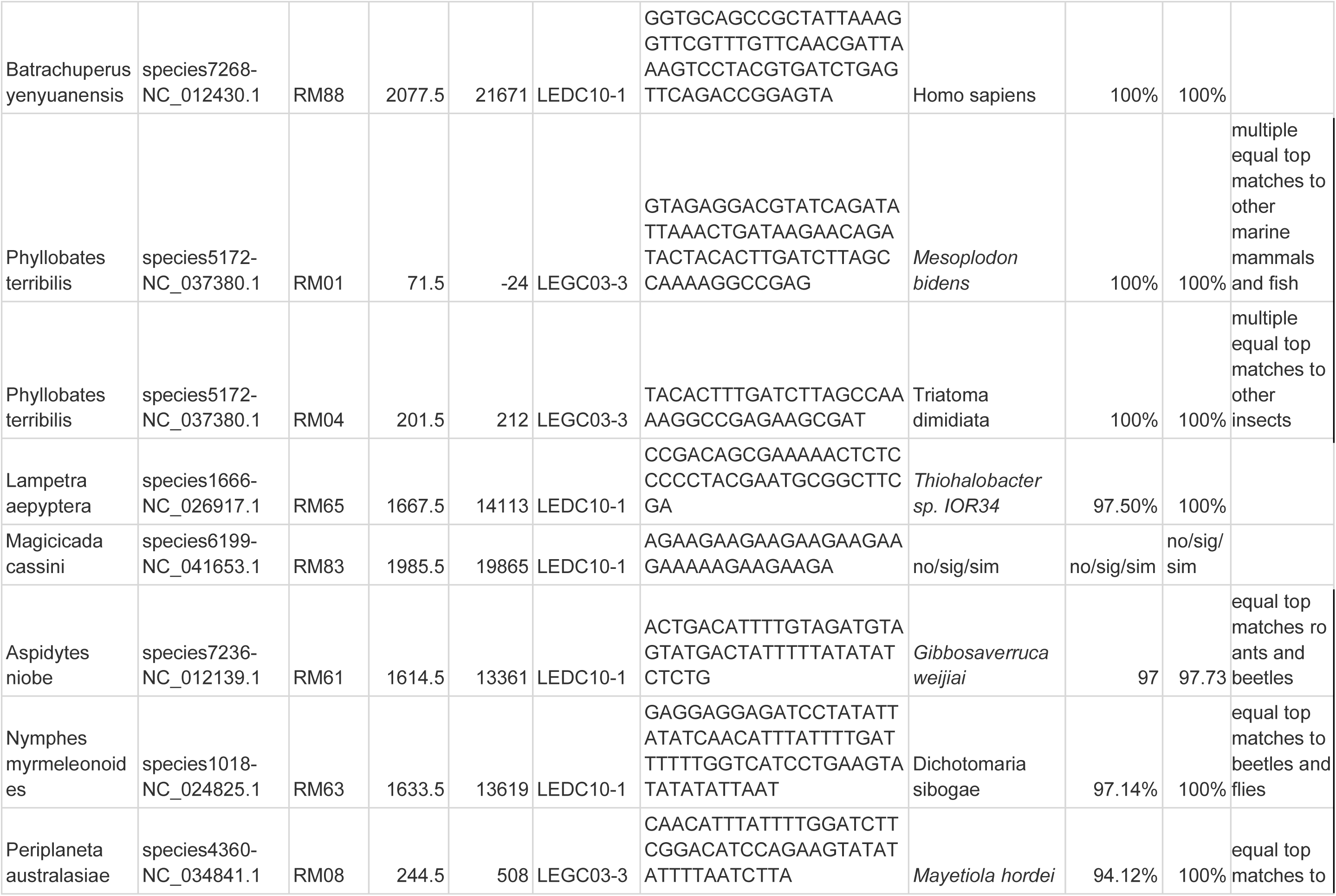
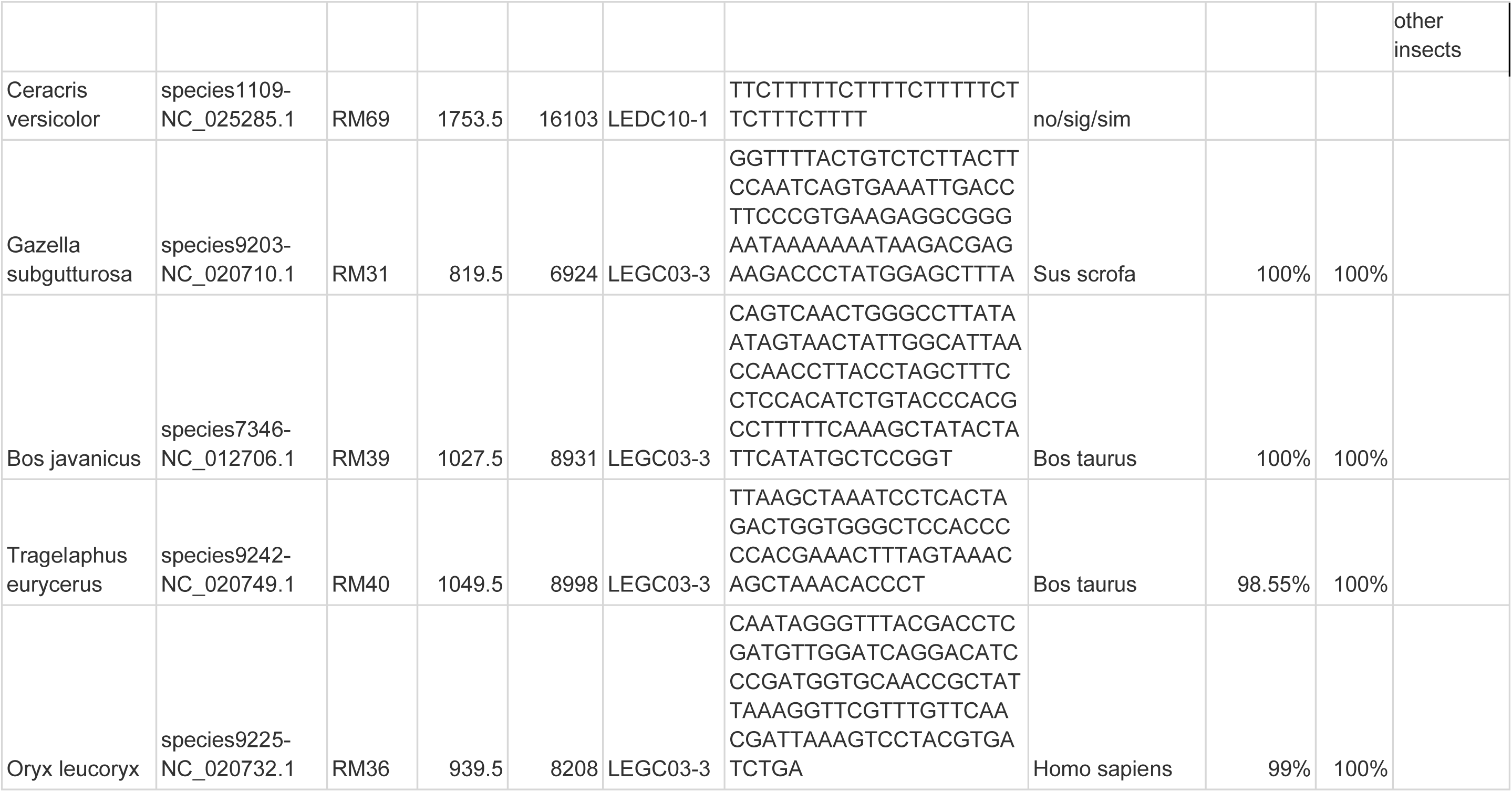
Blastn queries and results for animal and algal examples.

**Supplemental Table 3:** Each sample in the study has its carbon date in ybp. The youngest samples are nearly modern and, thus, have values of 0 or below. The following columns display the periods the sample age belongs to in three ways: human housing, paleoenvironmental, and fire time delineations.

| Sample | Core | Sample Depth | AGE | Housing Stage | Paleoclimate Events | Fire Stage |
| --- | --- | --- | --- | --- | --- | --- |
| RM01 | LEGC03-3 | 71.5 | -24 | European Arrival | Holocene | High-Fire |
| RM02 | LEGC03-3 | 99.5 | 0 | European Arrival | Holocene | High-Fire |
| RM03 | LEGC03-3 | 169.5 | 89 | European Arrival | Holocene | High-Fire |
| RM04 | LEGC03-3 | 201.5 | 212 | European Arrival | Holocene | High-Fire |
| RM05 | LEGC03-3 | 219.5 | 337 | European Arrival | Holocene | High-Fire |
| RM06 | LEGC03-3 | 229.5 | 403 | European Arrival | Holocene | High-Fire |
| RM08 | LEGC03-3 | 244.5 | 508 | Late Holocene | Holocene | High-Fire |
| RM09 | LEGC03-3 | 289.5 | 871 | Late Holocene | Holocene | High-Fire |
| RM10 | LEGC03-3 | 299.5 | 965 | Late Holocene | Holocene | High-Fire |
| RM11 | LEGC03-3 | 319.5 | 1433 | Late Holocene | Holocene | High-Fire |
| RM12 | LEGC03-3 | 349.5 | 2263 | Late Holocene | Holocene | High-Fire |
| RM13 | LEGC03-3 | 369.5 | 2346 | Late Holocene | Holocene | High-Fire |
| RM14 | LEGC03-3 | 384.5 | 2419 | Late Holocene | Holocene | High-Fire |
| RM15 | LEGC03-3 | 435.5 | 2698 | Late Holocene | Holocene | High-Fire |
| RM16 | LEGC03-3 | 447.5 | 2843 | Late Holocene | Holocene | High-Fire |
| RM17 | LEGC03-3 | 469.5 | 3084 | Late Holocene | Holocene | High-Fire |
| RM18 | LEGC03-3 | 489.5 | 3207 | Late Holocene | Holocene | High-Fire |
| RM19 | LEGC03-3 | 499.5 | 3273 | Late Holocene | Holocene | High-Fire |
| RM20 | LEGC03-3 | 529.5 | 3881 | Middle Holocene | Holocene | High-Fire |
| RM21 | LEGC03-3 | 579.5 | 4994 | Middle Holocene | Holocene | High-Fire |
| RM22 | LEGC03-3 | 599.5 | 5138 | Middle Holocene | Holocene | High-Fire |
| **RM23** | LEGC03-3 | 634.5 | 5463 | Middle<br>Holocene | Holocene | High-Fire |
| **RM25** | LEGC03-3 | 649.5 | 5591 | Middle<br>Holocene | Holocene | High-Fire |
| **RM26** | LEGC03-3 | 679.5 | 5862 | Middle<br>Holocene | Holocene | High-Fire |
| **RM27** | LEGC03-3 | 724.5 | 6363 | Middle<br>Holocene | Holocene | High-Fire |
| **RM28** | LEGC03-3 | 742.5 | 6496 | Middle<br>Holocene | Holocene | High-Fire |
| **RM29** | LEGC03-3 | 789.5 | 6719 | Middle<br>Holocene | Holocene | High-Fire |
| **RM30** | LEGC03-3 | 799.5 | 6758 | Middle<br>Holocene | Holocene | High-Fire |
| **RM31** | LEGC03-3 | 819.5 | 6924 | Middle<br>Holocene | Holocene | High-Fire |
| **RM32** | LEGC03-3 | 849.5 | 7401 | Early Holocene | Holocene | High-Fire |
| **RM33** | LEGC03-3 | 859.5 | 7702 | Early Holocene | Holocene | High-Fire |
| **RM34** | LEGC03-3 | 869.5 | 7763 | Early Holocene | Holocene | High-Fire |
| **RM35** | LEGC03-3 | 898.5 | 7931 | Early Holocene | Holocene | High-Fire |
| **RM36** | LEGC03-3 | 939.5 | 8208 | Early Holocene | Holocene | High-Fire |
| **RM37** | LEGC03-3 | 964.5 | 8614 | Early Holocene | Holocene | High-Fire |
| **RM38** | LEGC03-3 | 1019.5 | 8906 | Early Holocene | Holocene | High-Fire |
| **RM39** | LEGC03-3 | 1027.5 | 8931 | Early Holocene | Holocene | High-Fire |
| **RM40** | LEGC03-3 | 1049.5 | 8998 | Early Holocene | Holocene | High-Fire |
| **RM41** | LEGC03-3 | 1072.5 | 9065 | Early Holocene | Holocene | High-Fire |
| **RM42** | LEDC10-1 | 1084.5 | 9096 | Early Holocene | Holocene | High-Fire |
| **RM43** | LEDC10-1 | 1328.5 | 10135 | Early Holocene | Holocene | Intense-Fire |
| **RM44** | LEDC10-1 | 1338.5 | 10225 | Early Holocene | Holocene | Intense-Fire |
| **RM45** | LEDC10-1 | 1348.5 | 10319 | Early Holocene | Holocene | Intense-Fire |
| **RM46** | LEDC10-1 | 1358.5 | 10413 | Early Holocene | Holocene | Intense-Fire |
| **RM47** | LEDC10-1 | 1407.5 | 10848 | Population<br>Intensification | Holocene | Intense-Fire |
| **RM49** | LEDC10-1 | 1417.5 | 10942 | Population<br>Intensification | Holocene | Intense-Fire |
| **RM50** | LEDC10-1 | 1440.5 | 11184 | Population<br>Intensification | Holocene | Intense-Fire |
| **RM51** | LEDC10-1 | 1450.5 | 11327 | Population<br>Intensification | Holocene | Intense-Fire |
| **RM52** | LEDC10-1 | 1470.5 | 11612 | Population<br>Intensification | Holocene | Intense-Fire |
| **RM53** | LEDC10-1 | 1480.5 | 11753 | Population<br>Intensification | Younger Dryas | Intense-Fire |
| **RM54** | LEDC10-1 | 1511.5 | 12200 | Population<br>Intensification | Younger Dryas | Intense-Fire |
| **RM55** | LEDC10-1 | 1521.5 | 12387 | Population<br>Intensification | Younger Dryas | Intense-Fire |
| **RM56** | LEDC10-1 | 1531.5 | 12632 | Population<br>Intensification | Younger Dryas | Intense-Fire |
| **RM57** | LEDC10-1 | 1545.5 | 12860 | Population<br>Intensification | Bolling Allerod<br>Interstadial | Intense-Fire |
| **RM58** | LEDC10-1 | 1595.5 | 13224 | Debated<br>Presence | Bolling Allerod<br>Interstadial | Intense-Fire |
| **RM59** | LEDC10-1 | 1605.5 | 13296 | Debated<br>Presence | Bolling Allerod<br>Interstadial | Low-Fire |
| **RM61** | LEDC10-1 | 1614.5 | 13361 | Debated<br>Presence | Bolling Allerod<br>Interstadial | Low-Fire |
| **RM62** | LEDC10-1 | 1624.5 | 13479 | Debated<br>Presence | Bolling Allerod<br>Interstadial | Low-Fire |
| **RM63** | LEDC10-1 | 1633.5 | 13619 | Debated<br>Presence | Bolling Allerod<br>Interstadial | Low-Fire |
| **RM64** | LEDC10-1 | 1655.5 | 13948 | Debated<br>Presence | Bolling Allerod<br>Interstadial | Low-Fire |
| **RM65** | LEDC10-1 | 1667.5 | 14113 | Debated<br>Presence | Bolling Allerod<br>Interstadial | Low-Fire |
| **RM66** | LEDC10-1 | 1684.5 | 14355 | Debated<br>Presence | Bolling Allerod<br>Interstadial | Low-Fire |
| **RM67** | LEDC10-1 | 1717.5 | 15010 | Debated<br>Presence | HS-1 | Low-Fire |
| **RM68** | LEDC10-1 | 1729.5 | 15400 | Debated<br>Presence | HS-1 | Low-Fire |
| **RM69** | LEDC10-1 | 1753.5 | 16103 | Debated<br>Presence | HS-1 | Low-Fire |
| **RM70** | LEDC10-1 | 1764.5 | 16373 | Debated<br>Presence | HS-1 | Low-Fire |
| **RM71** | LEDC10-1 | 1785.5 | 16821 | Debated<br>Presence | HS-1 | Low-Fire |
| **RM74** | LEDC10-1 | 1803.5 | 17130 | Debated<br>Presence | HS-1 | Low-Fire |
| **RM75** | LEDC10-1 | 1816.5 | 17318 | Debated<br>Presence | HS-1 | Low-Fire |
| **RM76** | LEDC10-1 | 1845.5 | 17694 | Debated<br>Presence | HS-1 | Low-Fire |
| **RM77** | LEDC10-1 | 1860.5 | 17891 | Debated<br>Presence | HS-1 | Low-Fire |
| **RM78** | LEDC10-1 | 1875.5 | 18098 | Debated<br>Presence | HS-1 | Low-Fire |
| **RM79** | LEDC10-1 | 1897.5 | 18445 | Debated<br>Presence | HS-1 | Low-Fire |
| **RM80** | LEDC10-1 | 1936.5 | 19042 | Pre-Archaeological<br>Evidence | Last Glacial<br>Maximum | Low-Fire |
| **RM81** | LEDC10-1 | 1958.5 | 19474 | Pre-Archaeological<br>Evidence | Last Glacial<br>Maximum | Low-Fire |
| **RM82** | LEDC10-1 | 1968.5 | 19615 | Pre-Archaeological<br>Evidence | Last Glacial<br>Maximum | Low-Fire |
| **RM83** | LEDC10-1 | 1985.5 | 19865 | Pre-Archaeological<br>Evidence | Last Glacial<br>Maximum | Low-Fire |
| **RM84** | LEDC10-1 | 2012.5 | 20290 | Pre-Archaeological<br>Evidence | Last Glacial<br>Maximum | Low-Fire |
| **RM85** | LEDC10-1 | 2027.5 | 20535 | Pre-Archaeological Evidence | Last Glacial Maximum | Low-Fire |
| **RM86** | LEDC10-1 | 2045.5 | 20884 | Pre-Archaeological Evidence | Last Glacial Maximum | Low-Fire |
| **RM87** | LEDC10-1 | 2060.5 | 21242 | Pre-Archaeological Evidence | Last Glacial Maximum | Low-Fire |
| **RM88** | LEDC10-1 | 2077.5 | 21671 | Pre-Archaeological Evidence | Last Glacial Maximum | Low-Fire |
| **RM89** | LEDC10-1 | 2127.5 | 22466 | Pre-Archaeological Evidence | Last Glacial Maximum | Low-Fire |
| **RM90** | LEDC10-1 | 2139.5 | 22664 | Pre-Archaeological Evidence | Last Glacial Maximum | Low-Fire |
| **RM91** | LEDC10-1 | 2148.5 | 22821 | Pre-Archaeological Evidence | Last Glacial Maximum | Low-Fire |
| **RM92** | LEDC10-1 | 2172.5 | 23202 | Pre-Archaeological Evidence | Last Glacial Maximum | Low-Fire |
| **RM93** | LEDC10-1 | 2190.5 | 23491 | Pre-Archaeological Evidence | Last Glacial Maximum | Low-Fire |
| **RM94** | LEDC10-1 | 2213.5 | 23927 | Pre-Archaeological Evidence | Last Glacial Maximum | Low-Fire |

**Supplemental Table 4:** Genera within plant Orders presented in this paper that were likely to occur in the Lake Elsinore area. The list was made by using CalFlora data, selecting only native genera present in Riverside County. Genera were further filtered to include only those likely to appear in the habitat types or plant communities at Lake Elsinore and with records near Lake Elsinore. Observations made at Lake Elsinore were also included.

| **ORDER** | **FAMILY** | **GENERA** | **COMMUNITIES** |
| --- | --- | --- | --- |
| Alismatales | Alismataceae | Alisma | wetland-riparian |
| Alismatales | Alismataceae | Echinodorus | wetland-riparian |
| Alismatales | Alismataceae | Sagittaria | wetland-riparian |
| Alismatales | Araceae | Lemna | wetland-riparian |
| Alismatales | Araceae | Wolffia | wetland-riparian |
| Alismatales | Hydrocharitaceae | Najas | wetland-riparian |
| Alismatales | Juncaginaceae | Triglochin | wetland-riparian |
| Alismatales | Potamogetonaceae | Potamogeton | wetland-riparian |
| Alismatales | Potamogetonaceae | Stuckenia | wetland-riparian |
| Alismatales | Ruppiaaceae | Ruppia | wetland-riparian |
| Poales | Cyperaceae | Bolboschoenus | wetland-riparian |
| Poales | Cyperaceae | Carex | Foothill Woodland, Coastal Sage Scrub, Chaparral, Pinyon-Juniper Woodland, Valley grassland, wetland-riparian |
| Poales | Cyperaceae | Cyperus | Foothill Woodland, Coastal Sage Scrub, Southern Oak Woodland, wetland-riparian |
| Poales | Cyperaceae | Eleocharis | Coastal Sage Scrub, wetland-riparian |
| Poales | Cyperaceae | Isolepis | wetland-riparian |
| Poales | Cyperaceae | Schoenoplectus | wetland-riparian |
| Poales | Cyperaceae | Scirpus | Foothill Woodland, Chaparral, Valley grassland, wetland-riparian |
| Poales | Juncaceae | Juncus | Foothill Woodland, Chaparral, Coastal Sage Scrub, Valley Grassland, wetland-riparian |
| Poales | Poaceae | Agrostis | Foothill Woodland, Chaparral, Valley Grassland, wetland-riparian |
| Poales | Poaceae | Alopecurus | Coastal Sage Scrub, Chaparral, Valley Grassland, wetland-riparian |
| Poales | Poaceae | Andropogon | Coastal Sage Scrub, Chaparral, wetland-riparian |
| Poales | Poaceae | Aristida | Coastal Sage Scrub, Pinyon-Juniper Woodland, Valley Grassland |
| Poales | Poaceae | Bothriochloa | Coastal Sage Scrub |
| Poales | Poaceae | Bouteloua | Pinyon-Juniper Woodland |
| Poales | Poaceae | Bromus | Coastal Sage Scrub, Foothill Woodland, Chaparral, Valley Grassland, Pinyon-Juniper Woodland, wetland-riparian |
| Poales | Poaceae | Calamagrostis | Chaparral |
| Poales | Poaceae | Deschampsia | Coastal Sage Scrub, Foothill Woodland, Chaparral, Valley Grassland, wetland-riparian |
| Poales | Poaceae | Distichlis | Valley Grassland, wetland-riparian |
| Poales | Poaceae | Elymus | Coastal Sage Scrub, Southern Oak Woodland, Foothill Woodland, Chaparral, Valley Grassland, wetland-riparian |
| Poales | Poaceae | Eragrostis | wetland-riparian |
| Poales | Poaceae | Eriochloa | wetland-riparian |
| Poales | Poaceae | Festuca | Foothill Woodland, Chaparral, Valley Grassland, wetland-riparian |
| Poales | Poaceae | Hordeum | Foothill Woodland, Chaparral, Valley Grassland, wetland-riparian |
| Poales | Poaceae | Imperata | Coastal Sage Scrub, Chaparral, wetland-riparian |
| Poales | Poaceae | Koeleria | Foothill Woodland, Chaparral, Valley Grassland |
| Poales | Poaceae | Leptochloa | Valley Grassland, wetland-riparian |
| Poales | Poaceae | Melica | Coastal Sage Scrub, Foothill Woodland, Pinyon-Juniper Woodland |
| Poales | Poaceae | Muhlenbergia | Coastal Sage Scrub, Chaparral, Valley Grassland, wetland-riparian |
| Poales | Poaceae | Panicum | Valley Grassland, wetland-riparian |
| Poales | Poaceae | Paspalum | Valley Grassland, wetland-riparian |
| Poales | Poaceae | Phalaris | Coastal Sage Scrub, Foothill Woodland, Valley Grassland, wetland-riparian |
| Poales | Poaceae | Phragmites | Foothill Woodland, Chaparral, Valley Grassland, wetland-riparian |
| Poales | Poaceae | Poa | Foothill Woodland, Chaparral, Valley Grassland, Pinyon-Juniper Woodland |
| Poales | Poaceae | Scribneria | Foothill Woodland, Chaparral, Valley Grassland |
| Poales | Poaceae | Setaria | Coastal Sage Scrub, Valley Grassland |
| Poales | Poaceae | Sporobolus | Coastal Sage Scrub, Pinyon-Juniper Woodland, wetland-riparian |
| Poales | Poaceae | Stipa | Coastal Sage Scrub, Foothill Woodland, Chaparral, Pinyon-Juniper Woodland |
| Poales | Typhaceae | Sparganium | wetland-riparian |
| Poales | Typhaceae | Typha | wetland-riparian |
| Fagales | Betulaceae | Alnus | Foothill Woodland, Chaparral, wetland-riparian |
| Fagales | Fagaceae | Quercus | Coastal Sage Scrub, Chaparral, Southern Oak Woodland, Foothill Woodland, Valley Grassland, Pinyon-Juniper Woodland |
| Fagales | Juglandaceae | Juglans | Southern Oak Woodland, Foothill Woodland, wetland-riparian |
| Fabales | Fabaceae | Acmispon | Coastal Sage Scrub, Foothill Woodland, Chaparral, Pinyon-Juniper Woodland, Valley Grassland |
| Fabales | Fabaceae | Amorpha | Coastal Sage Scrub, Chaparral, wetland-riparian |
| Fabales | Fabaceae | Astragalus | Coastal Sage Scrub, Southern Oak Woodland, Foothill Woodland, Chaparral, Valley Grassland, Pinyon-Juniper Woodland, wetland-riparian |
| Fabales | Fabaceae | Glycyrrhiza | Foothill Woodland, Chaparral, Valley Grassland, wetland-riparian |
| Fabales | Fabaceae | Hoita | Foothill Woodland, Chaparral, Valley Grassland, wetland-riparian |
| Fabales | Fabaceae | Hosackia | wetland-riparian |
| Fabales | Fabaceae | Lupinus | Coastal Sage Scrub, Foothill Woodland, Chaparral, Pinyon-Juniper Woodland, Southern Oak Woodland, Valley Grassland, wetland-riparian |
| Fabales | Fabaceae | Lathyrus | Coastal Sage Scrub, Chaparral, Foothill Woodland |
| Fabales | Fabaceae | Neltuma | *Specific community information unavailable* |
| Fabales | Fabaceae | Pickeringia | Chaparral |
| Fabales | Fabaceae | Rupertia | Foothill Woodland, Chaparral |
| Fabales | Fabaceae | Thermopsis | Foothill Woodland, Valley Grassland |
| Fabales | Fabaceae | Trifolium | Coastal Sage Scrub, Foothill Woodland, Chaparral, Valley Grassland, wetland-riparian |
| Fabales | Fabaceae | Vicia | *Specific community information unavailable* |
| Fabales | Polygalaceae | Polygala | Pinyon-Juniper Woodland, Foothill Woodland, Chaparral, wetland-riparian |
| Fabales | Polygalaceae | Rhinotropis | *Specific community information unavailable* |
| Myrtales | Lythraceae | Ammannia | wetland-riparian |
| Myrtales | Lythraceae | Lythrum | Foothill Woodland, Chaparral, Valley Grassland, wetland-riparian |
| Myrtales | Onagraceae | Camissonia | Pinyon-Juniper Woodland |
| Myrtales | Onagraceae | Camissoniopsis | Coastal Sage Scrub, Southern Oak Woodland, Chaparral, Foothill Woodland, Valley Grassland |
| Myrtales | Onagraceae | Clarkia | Coastal Sage Scrub, Southern Oak Woodland, Chaparral, Foothill Woodland, Valley Grassland |
| Myrtales | Onagraceae | Epilobium | Coastal Sage Scrub, Foothill Woodland, Chaparral, Valley Grassland, Pinyon-Juniper Woodland, wetland-riparian |
| Myrtales | Onagraceae | Eulobus | Coastal Sage Scrub, Foothill Woodland, Chaparral, Valley Grassland |
| Myrtales | Onagraceae | Gayophytum | Chaparral, Pinyon-Juniper Woodland, wetland-riparian |
| Myrtales | Onagraceae | Tetrapteron | Foothill Woodland, Valley Grassland |
| Caryophyllales | Aizoaceae | Sesuvium | Coastal Sage Scrub, Valley Grassland, wetland-riparian |
| Caryophyllales | Amaranthaceae | Amaranthus | wetland-riparian |
| Caryophyllales | Amaranthaceae | Nitrophila | Foothill Woodland, Chaparral, Valley Grassland, wetland-riparian |
| Caryophyllales | Cactaceae | Cylindropuntia | Coastal Sage Scrub, Pinyon-Juniper Woodland |
| Caryophyllales | Cactaceae | Opuntia | Coastal Sage Scrub, Southern Oak Woodland, Chaparral, Valley Grassland, Pinyon-Juniper Woodland |
| Caryophyllales | Caryophyllaceae | Loeflingia | Coastal Sage Scrub, Chaparral, Valley Grassland |
| Caryophyllales | Caryophyllaceae | Polycarpon | Coastal Sage Scrub, Chaparral |
| Caryophyllales | Caryophyllaceae | Sabulina | *Specific community information unavailable* |
| Caryophyllales | Caryophyllaceae | Sagina | wetland-riparian |
| Caryophyllales | Caryophyllaceae | Silene | Coastal Sage Scrub, Chaparral |
| Caryophyllales | Caryophyllaceae | Spergularia | Coastal Sage Scrub, wetland-riparian |
| Caryophyllales | Portulacaceae | Calandrinia | Coastal Sage Scrub, Chaparral |
| Caryophyllales | Portulacaceae | Calyptridium | Coastal Sage Scrub, Foothill Woodland, Chaparral, Pinyon-Juniper Woodland |
| Caryophyllales | Portulacaceae | Claytonia | Coastal Sage Scrub, Foothill Woodland, Chaparral, Valley Grassland, Pinyon-Juniper Woodland |
| Caryophyllales | Portulacaceae | Montia | wetland-riparian |
| Caryophyllales | Nyctaginaceae | Abronia | Pinyon-Juniper Woodland |
| Caryophyllales | Nyctaginaceae | Boerhavia | Southern Oak Woodland, Valley Grassland |
| Caryophyllales | Nyctaginaceae | Mirabilis | Coastal Sage Scrub, Foothill Woodland, Chaparral, Pinyon-Juniper Woodland |
| Caryophyllales | Polygonaceae | Chorizanthe | Coastal Sage Scrub, Foothill Woodland, Chaparral, Valley Grassland, Pinyon-Juniper Woodland |
| Caryophyllales | Polygonaceae | Dodecahema | Coastal Sage Scrub, Chaparral |
| Caryophyllales | Polygonaceae | Eriogonum | Coastal Sage Scrub, Southern Oak Woodland, Foothill Woodland, Chaparral, Valley Grassland, Pinyon-Juniper Woodland |
| Caryophyllales | Polygonaceae | Lastarriaea | Coastal Sage Scrub, Chaparral |
| Caryophyllales | Polygonaceae | Persicaria | wetland-riparian |
| Caryophyllales | Polygonaceae | Polygonum | wetland-riparian |
| Crossosomatales | Crossosomataceae | Glossopetalon | *Specific community information unavailable* |
| Crossosomatales | Crossosomataceae | Crossosoma | *Specific community information unavailable* |
| Brassicales | Brassicaceae | Athysanus | Foothill Woodland, Chaparral, Valley Grassland |
| Brassicales | Brassicaceae | Barbarea | wetland-riparian |
| Brassicales | Brassicaceae | Cardamine | Southern Oak Woodland, Foothill Woodland, Chaparral, wetland-riparian |
| Brassicales | Brassicaceae | Descurainia | Foothill Woodland, Chaparral, Valley Grassland, Pinyon-Juniper Woodland |
| Brassicales | Brassicaceae | Draba | Coastal Sage Scrub, Chaparral, Pinyon-Juniper Woodland |
| Brassicales | Brassicaceae | Erysimum | Coastal Sage Scrub, Foothill Woodland, Chaparral, Valley Grassland, Pinyon-Juniper Woodland |
| Brassicales | Brassicaceae | Lepidium | Coastal Sage Scrub, Chaparral, Valley Grassland, Pinyon-Juniper Woodland, wetland-riparian |
| Brassicales | Brassicaceae | Nasturtium | wetland-riparian |
| Brassicales | Brassicaceae | Planodes | Coastal Sage Scrub, Valley Grassland, wetland-riparian |
| Brassicales | Brassicaceae | Rorippa | Foothill Woodland, Chaparral, Valley Grassland, wetland-riparian |
| Brassicales | Brassicaceae | Sibaropsis | *Specific community information unavailable* |
| Brassicales | Brassicaceae | Thysanocarpus | Coastal Sage Scrub, Foothill Woodland, Chaparral, Valley Grassland, Pinyon-Juniper Woodland |
| Brassicales | Brassicaceae | Tropidocarpum | Foothill Woodland, Valley Grassland |
| Brassicales | Brassicaceae | Turritis | Foothill Woodland, Chaparral, Valley Grassland |
| Brassicales | Cleomaceae | Cleomella | *Specific community information unavailable* |
| Brassicales | Limnanthaceae | Limnanthes | Foothill Woodland, Valley Grassland, wetland-riparian |
| Brassicales | Resedaceae | Oligomeris | *Specific community information unavailable* |
| Malvales | Malvaceae | Ayenia | *Specific community information unavailable* |
| Malvales | Malvaceae | Eremalche | *Specific community information unavailable* |
| Malvales | Malvaceae | Fremontodendron | Chaparral, Pinyon-Juniper Woodland |
| Malvales | Malvaceae | Malacothamnus | Coastal Sage Scrub, Foothill Woodland, Chaparral |
| Malvales | Malvaceae | Malvella | wetland-riparian |
| Malvales | Malvaceae | Sidalcea | Coastal Sage Scrub, Foothill Woodland, Valley Grassland, Chaparral, wetland-riparian |
| Malvales | Malvaceae | Sphaeralcea | Coastal Sage Scrub, Pinyon-Juniper Woodland |
| Malvales | Cistaceae | Crocanthemum | Chaparral |
| Lamiales | Bignoniaceae | Chilopsis | wetland-riparian |
| Lamiales | Boraginaceae | Amsinckia | Foothill Woodland, Chaparral, Valley Grassland |
| Lamiales | Boraginaceae | Cryptantha | Coastal Sage Scrub, Foothill Woodland, Chaparral, Valley Grassland, Pinyon-Juniper Woodland |
| Lamiales | Boraginaceae | Harpagonella | Coastal Sage Scrub, Chaparral, Valley Grassland |
| Lamiales | Boraginaceae | Pectocarya | Coastal Sage Scrub, Foothill Woodland, Chaparral, Valley Grassland, Pinyon-Juniper Woodland |
| Lamiales | Boraginaceae | Plagiobothrys | Coastal Sage Scrub, Foothill Woodland, Chaparral, Valley Grassland, Pinyon-Juniper Woodland, wetland-riparian |
| Lamiales | Lamiaceae | Clinopodium | Coastal Sage Scrub, Foothill Woodland, Chaparral, Valley Grassland |
| Lamiales | Lamiaceae | Lepechinia | Foothill Woodland, Chaparral |
| Lamiales | Lamiaceae | Monardella | Coastal Sage Scrub, Foothill Woodland, Chaparral, Pinyon-Juniper Woodland |
| Lamiales | Lamiaceae | Pycnanthemum | Foothill Woodland, Chaparral |
| Lamiales | Lamiaceae | Salvia | Coastal Sage Scrub, Foothill Woodland, Chaparral, Valley Grassland, Pinyon-Juniper Woodland |
| Lamiales | Lamiaceae | Scutellaria | Foothill Woodland, Chaparral, wetland-riparian |
| Lamiales | Lamiaceae | Stachys | Coastal Sage Scrub, Foothill Woodland, Chaparral, wetland-riparian |
| Lamiales | Lamiaceae | Trichostema | Coastal Sage Scrub, Southern Oak Woodland, Foothill Woodland, Chaparral, wetland-riparian |
| Lamiales | Oleaceae | Forestiera | Coastal Sage Scrub, Foothill Woodland, Chaparral |
| Lamiales | Oleaceae | Fraxinus | Southern Oak Woodland, Foothill Woodland, Chaparral, Valley Grassland, wetland-riparian |
| Lamiales | Orobanchaceae | Aphyllon | Chaparral, Pinyon-Juniper Woodland |
| Lamiales | Orobanchaceae | Castilleja | Coastal Sage Scrub, Foothill Woodland, Chaparral, Valley Grassland, Pinyon-Juniper Woodland, wetland-riparian |
| Lamiales | Orobanchaceae | Cordylanthus | Foothill Woodland, Chaparral |
| Lamiales | Orobanchaceae | Pedicularis | Foothill Woodland, Chaparral |
| Lamiales | Phymaceae | Diplacus | Coastal Sage Scrub, Foothill Woodland, Chaparral, Pinyon-Juniper Woodland |
| Lamiales | Phymaceae | Mimetanthe | Foothill Woodland, Chaparral, Valley Grassland, wetland-riparian |
| Lamiales | Plantaginaceae | Antirrhinum | Coastal Sage Scrub, Chaparral |
| Lamiales | Plantaginaceae | Callitriche | Foothill Woodland, Chaparral, Valley Grassland, wetland-riparian |
| Lamiales | Plantaginaceae | Keckiella | Chaparral, Pinyon-Juniper Woodland |
| Lamiales | Plantaginaceae | Nuttallanthus | Foothill Woodland, Chaparral, Valley Grassland |
| Lamiales | Plantaginaceae | Penstemon | Coastal Sage Scrub, Foothill Woodland, Chaparral, Pinyon-Juniper Woodland |
| Lamiales | Plantaginaceae | Plantago | Coastal Sage Scrub, Foothill Woodland, Chaparral, Pinyon-Juniper Woodland, wetland-riparian |
| Lamiales | Plantaginaceae | Veronica | Foothill Woodland, Chaparral, Valley Grassland, wetland-riparian |
| Lamiales | Scrophulariaceae | Scrophularia | Coastal Sage Scrub Foothill Woodland, Chaparral, Valley Grassland, wetland-riparian |
| Lamiales | Verbenaceae | Phyla | Valley Grassland, wetland-riparian |
| Lamiales | Verbenaceae | Verbena | Foothill Woodland, Chaparral, Valley Grassland, Pinyon-Juniper Woodland, wetland-riparian |
| Gentianales | Apocynaceae | Apocynum | Foothill Woodland, Chaparral, Valley Grassland, wetland-riparian |
| Gentianales | Apocynaceae | Asclepias | Foothill Woodland, Chaparral, Valley Grassland, Pinyon-Juniper Woodland, wetland-riparian |
| Gentianales | Apocynaceae | Funastrum | Coastal Sage Scrub, Chaparral, wetland-riparian |
| Gentianales | Gentianaceae | Eustoma | Coastal Sage Scrub, wetland-riparian |
| Gentianales | Gentianaceae | Frasera | Coastal Sage Scrub, Southern Oak Woodland, Chaparral, Pinyon-Juniper Woodland |
| Gentianales | Gentianaceae | Zeltnera | Coastal Sage Scrub, Foothill Woodland, Chaparral, Valley Grassland, wetland-riparian |
| Gentianales | Rubiaceae | Galium | Foothill Woodland, Chaparral, Valley Grassland, Pinyon-Juniper Woodland, wetland-riparian |
| Rosales | Rosaceae | Adenostoma | Chaparral |
| Rosales | Rosaceae | Aphanes | Foothill Woodland, Valley Grassland |
| Rosales | Rosaceae | Cercocarpus | Chaparral, Pinyon-Juniper Woodland |
| Rosales | Rosaceae | Drymocallis | wetland-riparian |
| Rosales | Rosaceae | Heteromeles | Chaparral |
| Rosales | Rosaceae | Holodiscus | Chaparral, wetland-riparian |
| Rosales | Rosaceae | Horkelia | Coastal Sage Scrub, Foothill Woodland, Chaparral, Valley Grassland, wetland-riparian |
| Rosales | Rosaceae | Prunus | Coastal Sage Scrub, Foothill Woodland, Chaparral, Pinyon-Juniper Woodland, wetland-riparian |
| Rosales | Rosaceae | Rosa | Foothill Woodland, Chaparral, Valley Grassland, Pinyon-Juniper Woodland, wetland-riparian |
| Rosales | Rosaceae | Rubus | Foothill Woodland, Chaparral, Valley Grassland, wetland-riparian |
| Rosales | Urticaceae | Hesperocnide | *Specific community information unavailable* |
| Rosales | Urticaceae | Parietaria | wetland-riparian |
| Rosales | Urticaceae | Urtica | Foothill Woodland, Chaparral, Valley Grassland, wetland-riparian |
| Rosales | Rhamnaceae | Ceanothus | Foothill Woodland, Chaparral, Pinyon-Juniper Woodland |
| Rosales | Rhamnaceae | Frangula | Coastal Sage Scrub, Foothill Woodland, Chaparral, Pinyon-Juniper Woodland |
| Rosales | Rhamnaceae | Rhamnus | Coastal Sage Scrub, Southern Oak Woodland, Foothill Woodland, Chaparral |
| Asparagales | Iridaceae | Sisyrinchium | Foothill Woodland, wetland-riparian |
| Asparagales | Orchidaceae | Epipactis | Foothill Woodland, Chaparral, Valley Grassland, wetland-riparian |
| Asparagales | Orchidaceae | Piperia | Southern Oak Woodland, Foothill Woodland, Chaparral, Valley Grassland, wetland-riparian |
| Asparagales | Agavaceae | Yucca | Chaparral |
| Asparagales | Agavaceae | Chlorogalum | Coastal Sage Scrub, Foothill Woodland, Chaparral, Valley Grassland |
| Asparagales | Agavaceae | Hesperoyucca | Coastal Sage Scrub, Chaparral, Pinyon-Juniper Woodland |
| Asparagales | Agavaceae | Hooveria | *Specific community information unavailable* |
| Asparagales | Alliaceae | Allium | Coastal Sage Scrub, Southern Oak Woodland, Chaparral, Valley Grassland, Foothill Woodland, Pinyon-Juniper Woodland |

**Supplemental Figure 1:**
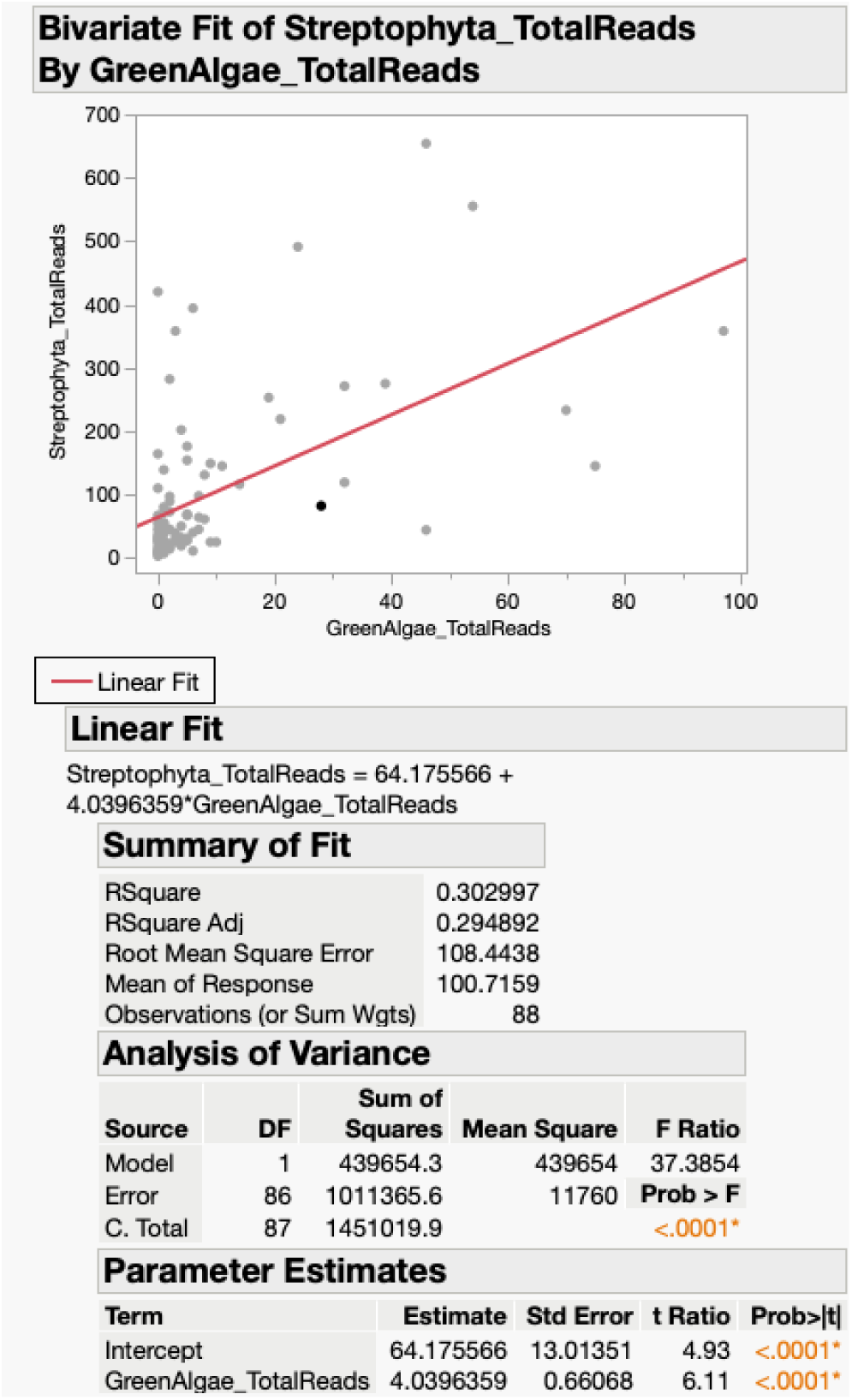
Bivariate plot of the per-sample abundance of algae and land plants. The Y axis represents the total Streptophyta reads per sample, while the X axis represents the total Chlorophyta reads per sample. The positive linear relationship between the two is significant (p < 0.001).

**Supplemental Figure 2:**
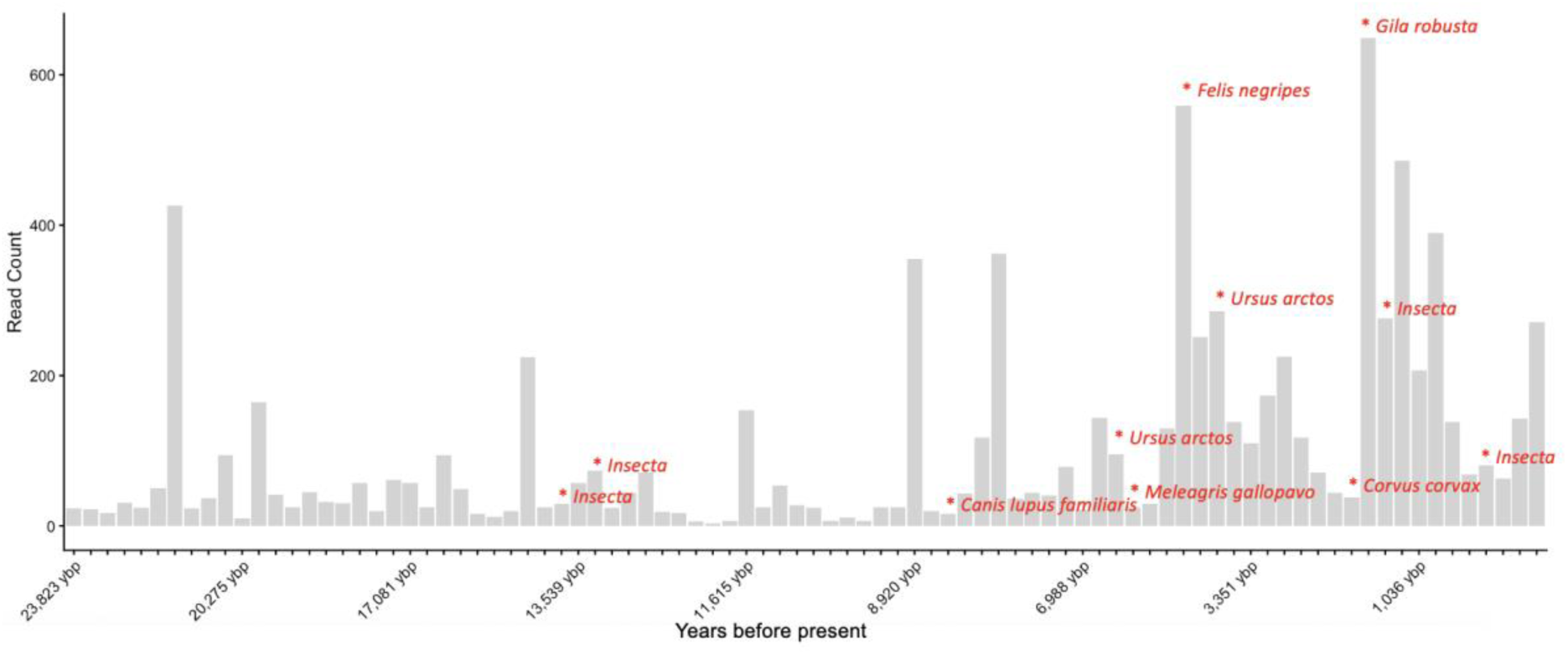
Counts of summed Streptophyta reads per sample. Taxon names in red are the top BLAST hit. While unique animal sequences were not in high enough in abundance per species to verify characteristic damage indicative of ancientness with mapDamage, some individual BLAST queries showed damage patterns (Supplementary Table 2).

**Supplemental Figure 3:**
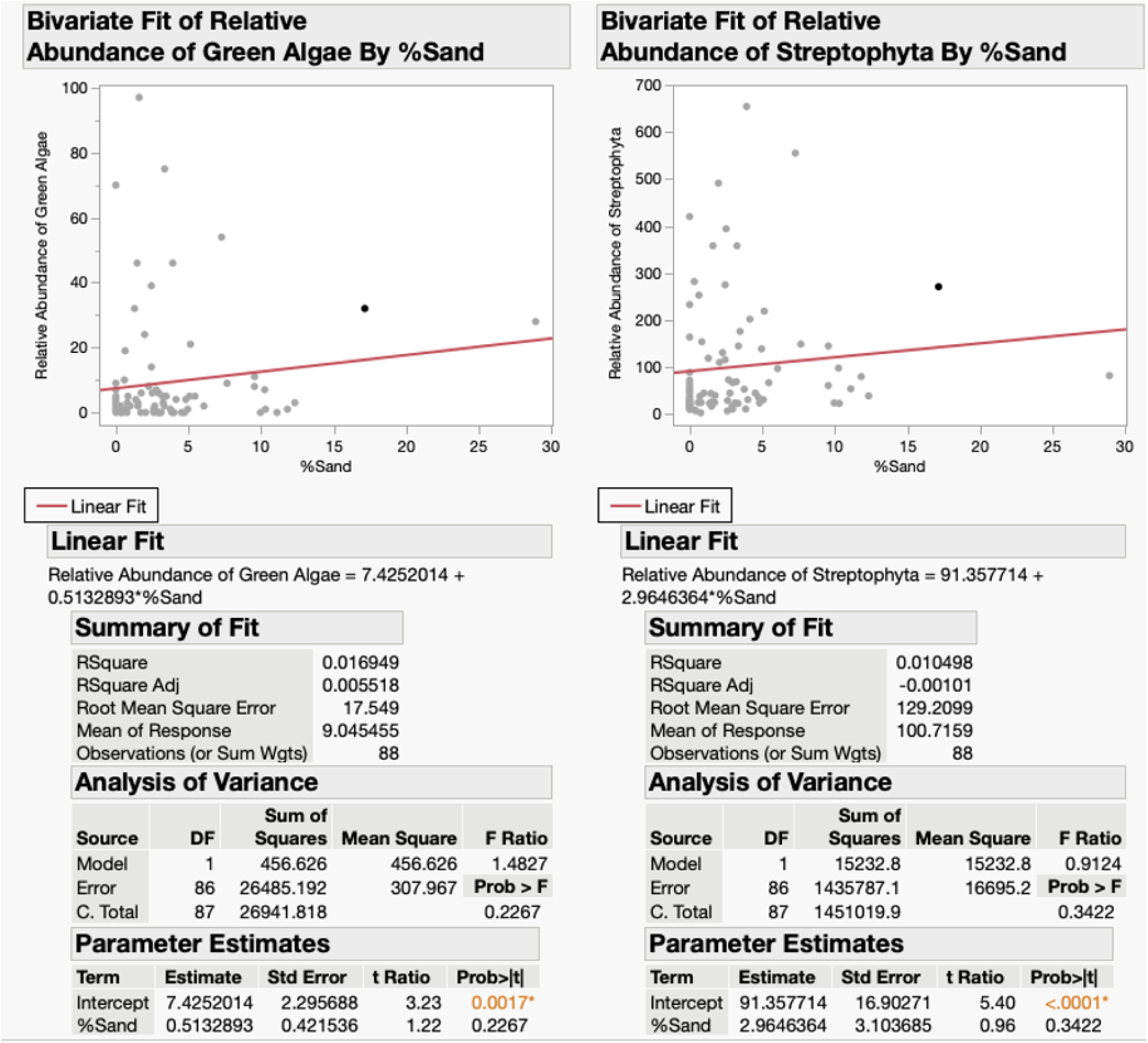
Bivariate plots of per sample abundance of algae (left) and land plants (right) against percent sand. Both linear regressions are insignificant (p > 0.05)

**Supplemental Figure 4:**
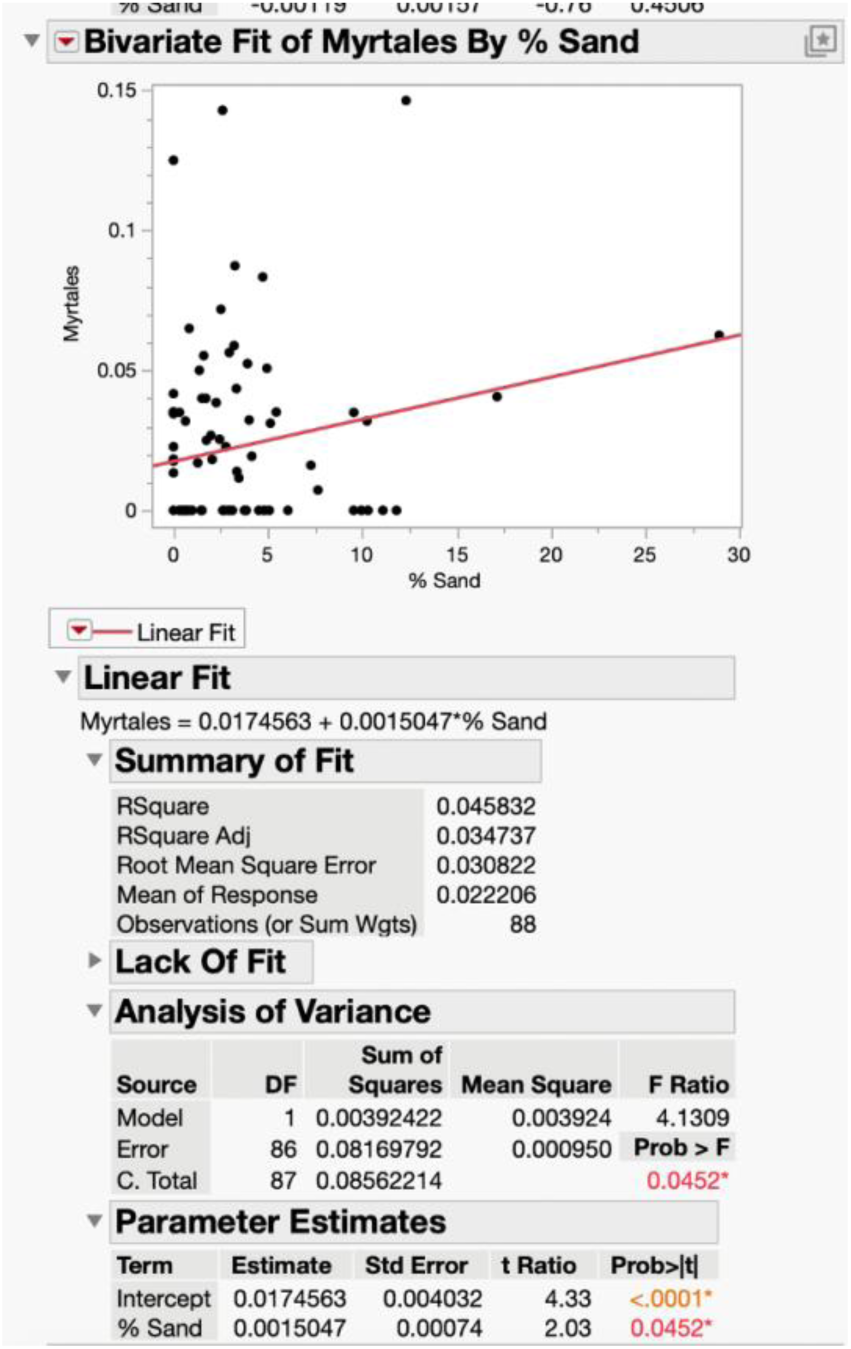
Linear regressions of relative abundance of relative abundance of Myrtales vs % Sand.

**Supplemental Figure 5:**
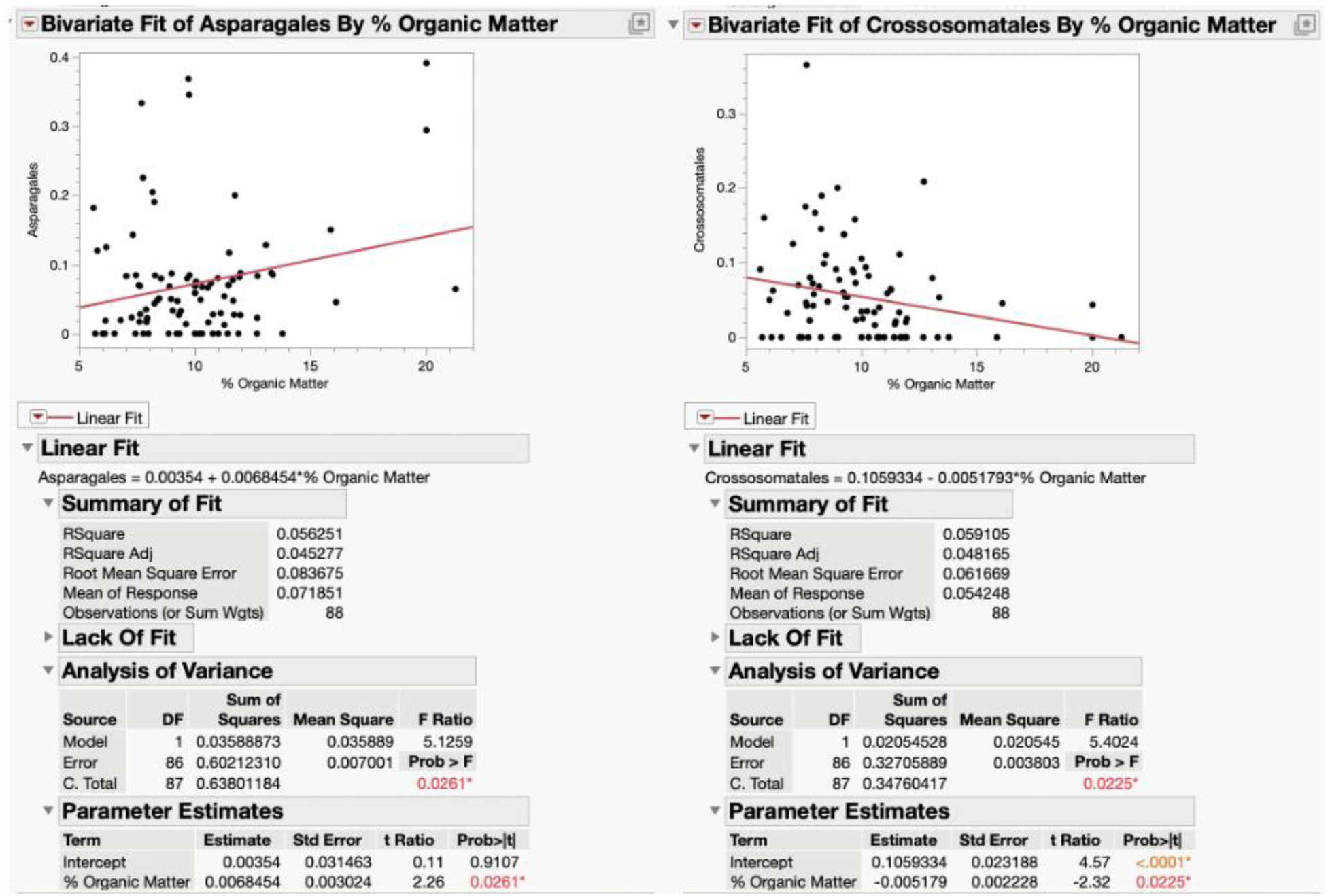
Linear regressions of relative abundance of Crossosomatales and Asparagales vs % Organic Matter.

**Supplemental Document 1**; Compilation of fragment misincorporation plots from multiple species generated by mapDamage.

**Supplemental Document 2**; Spreadsheet of Kruskall-Wallis and Dunn Test results.

**Supplemental Document 3**; Dataframe of the Target Capture data used for analyses that has been filtered with a prevalence filter of 20%, has blank samples removed, and removed targets with singleton reads.

