## Supplementary figures and images for "First Plant sedaDNA assemblage from California reveals ecological stability throughout 10,000 years of human presence at Lake Elsinore"

### Supplemental Document 1

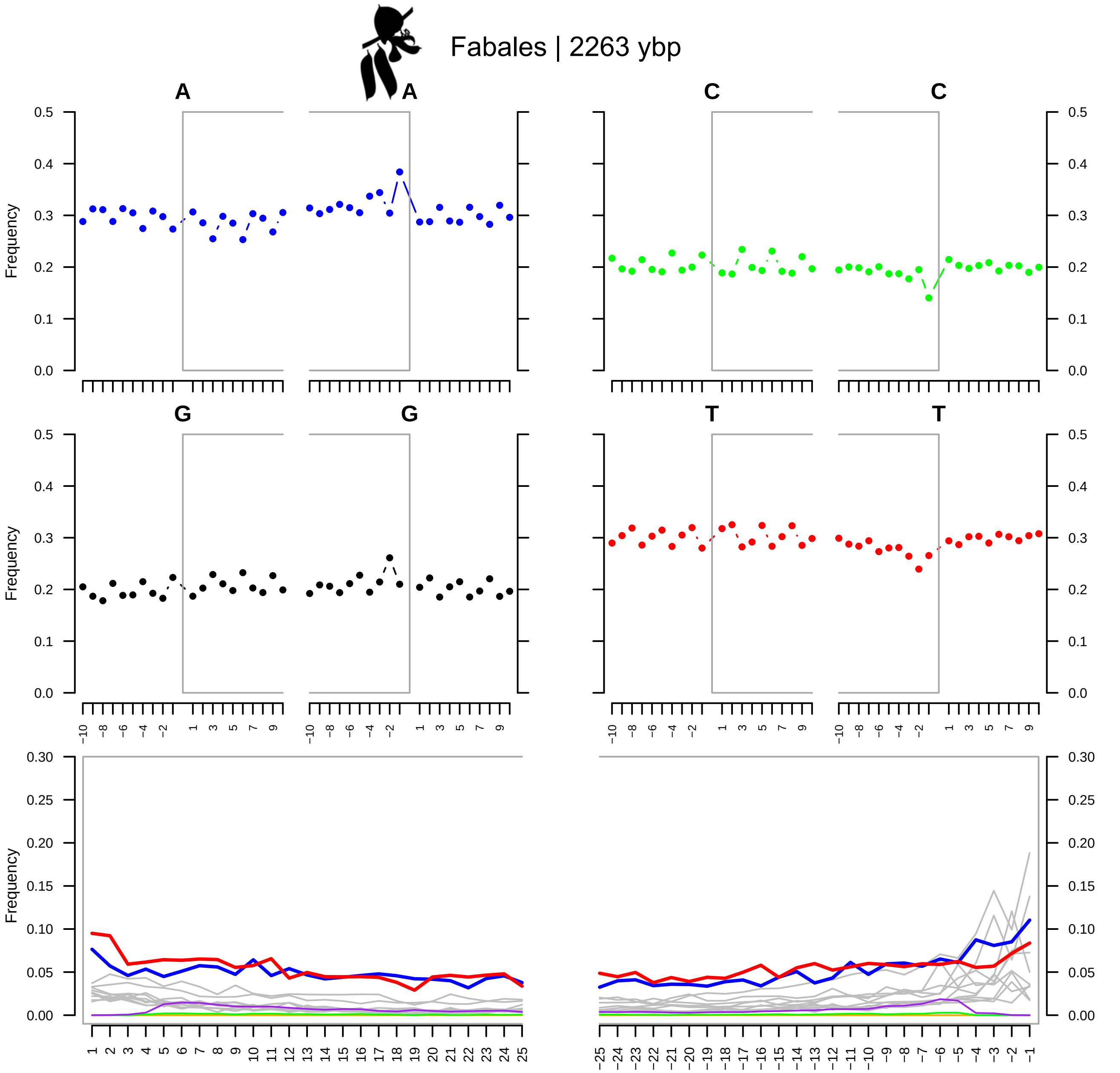

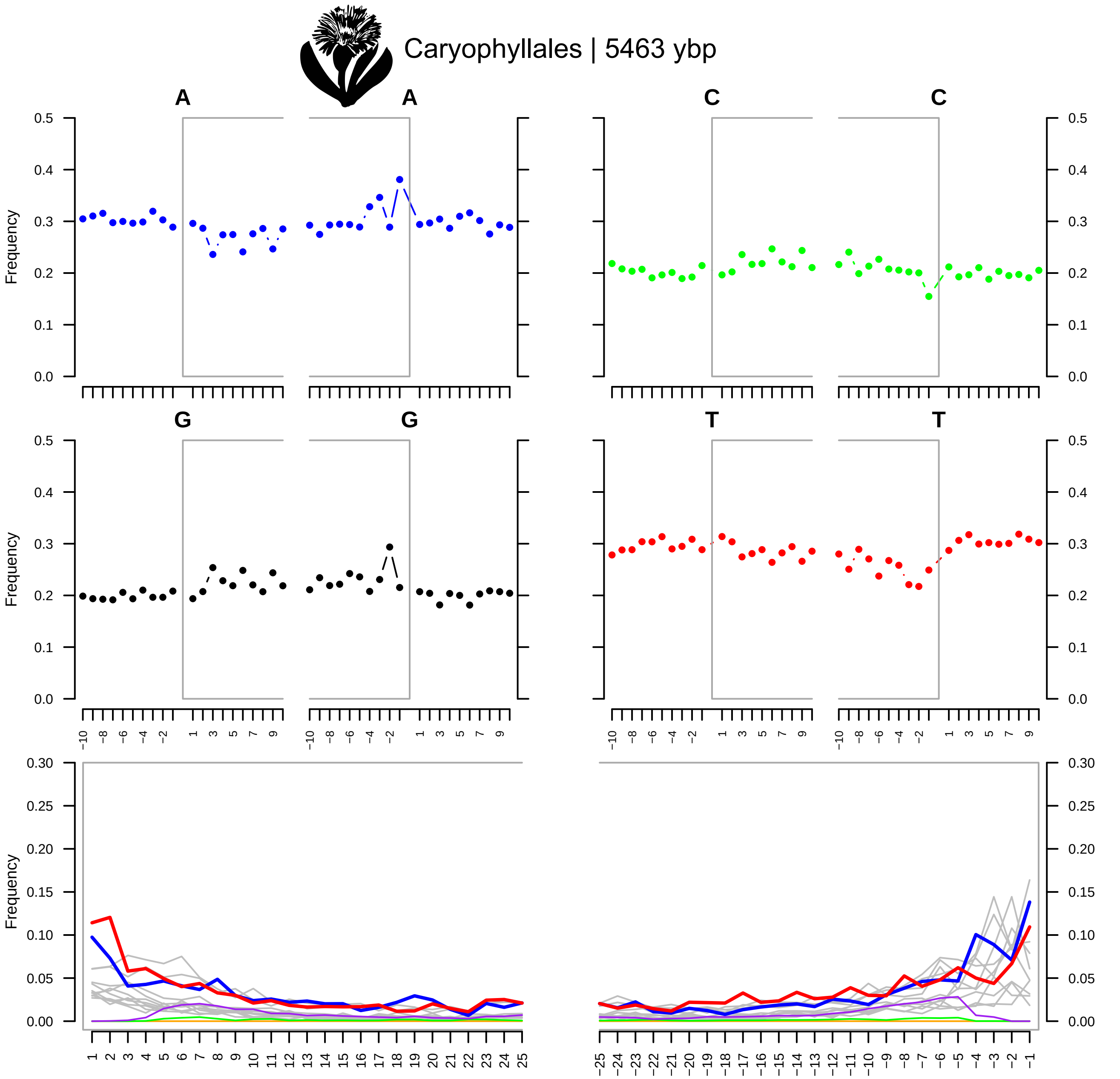

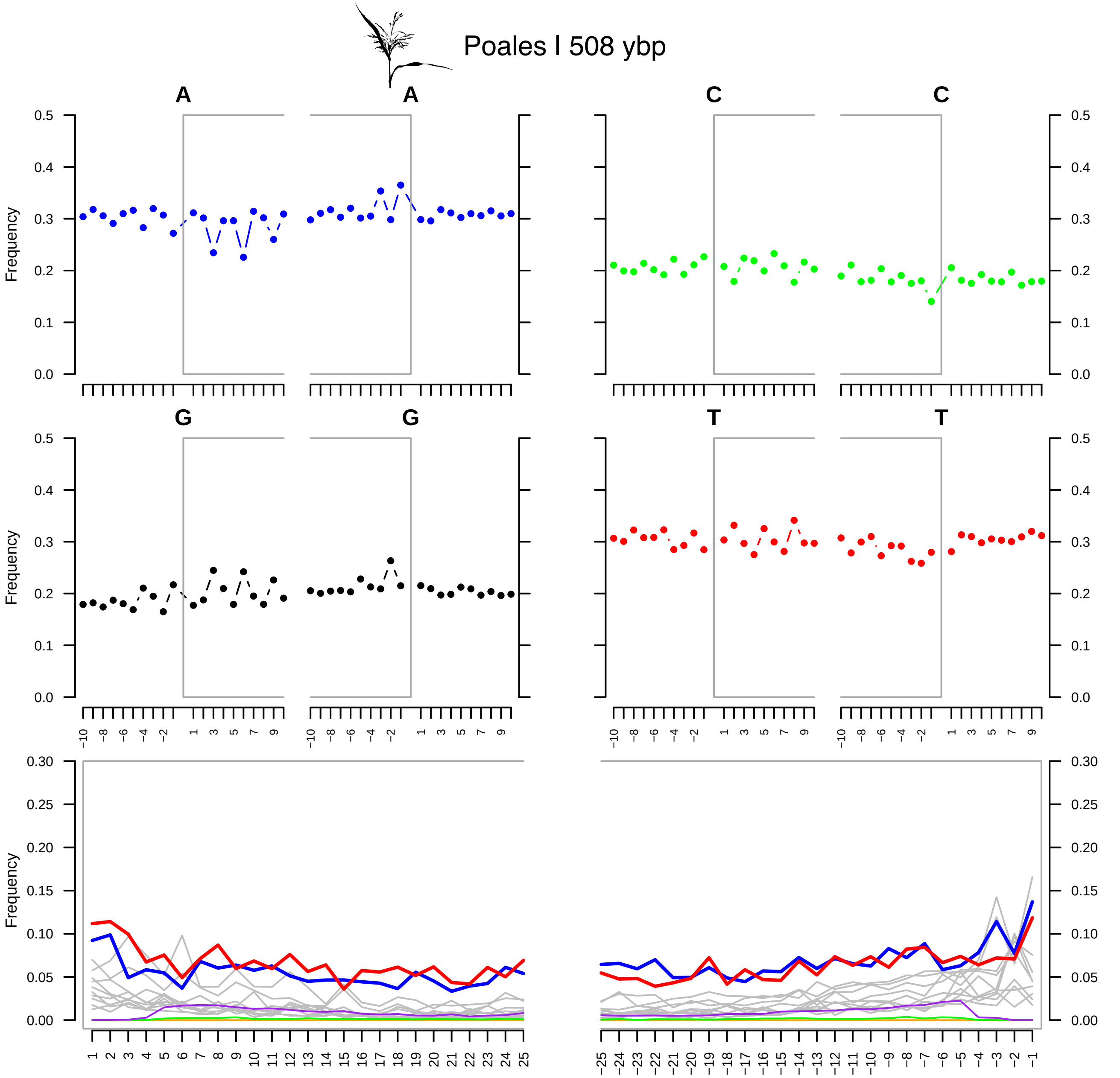

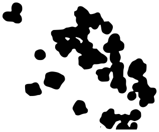

*Nannochloropsis limnetica* | 7931

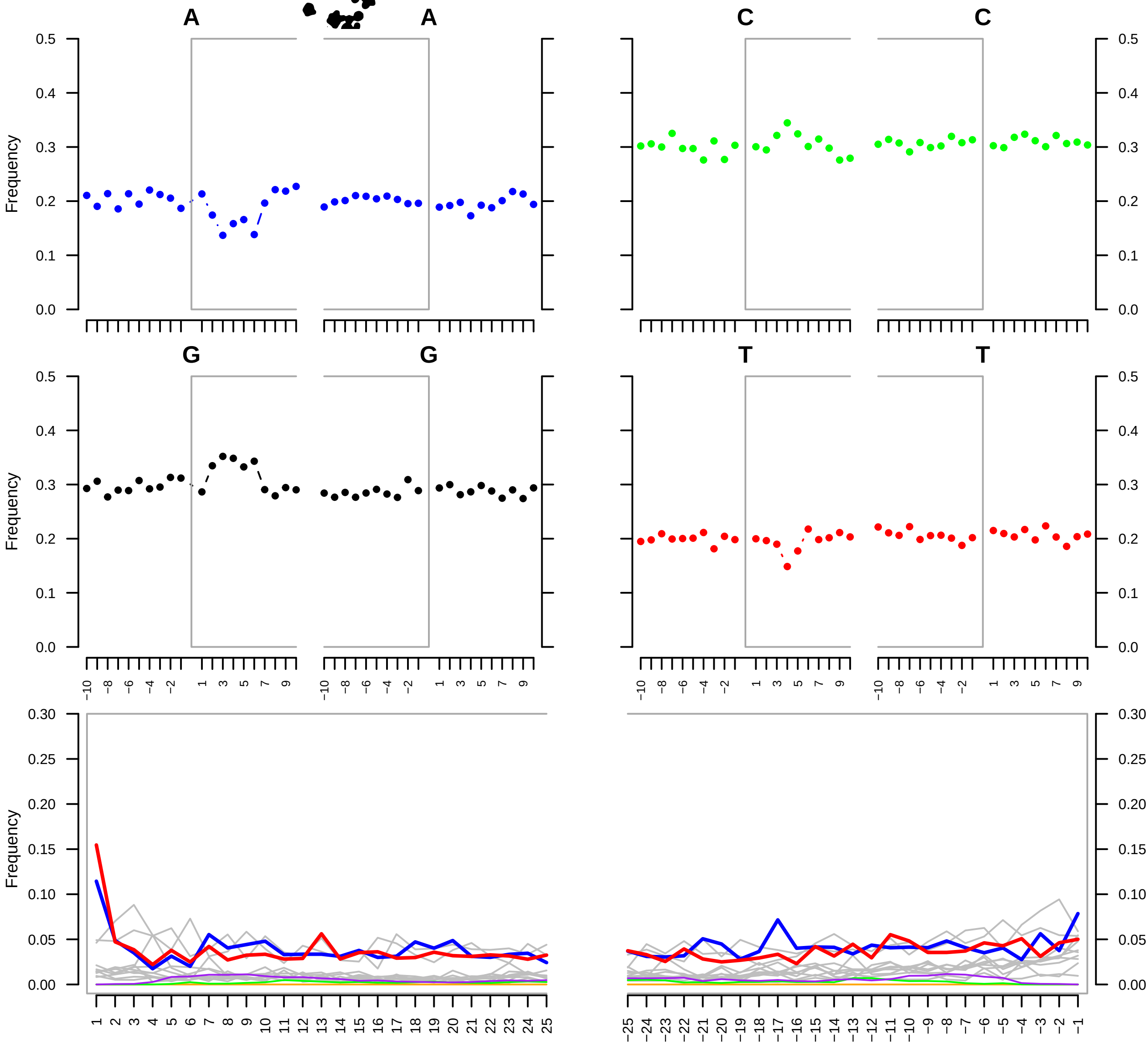

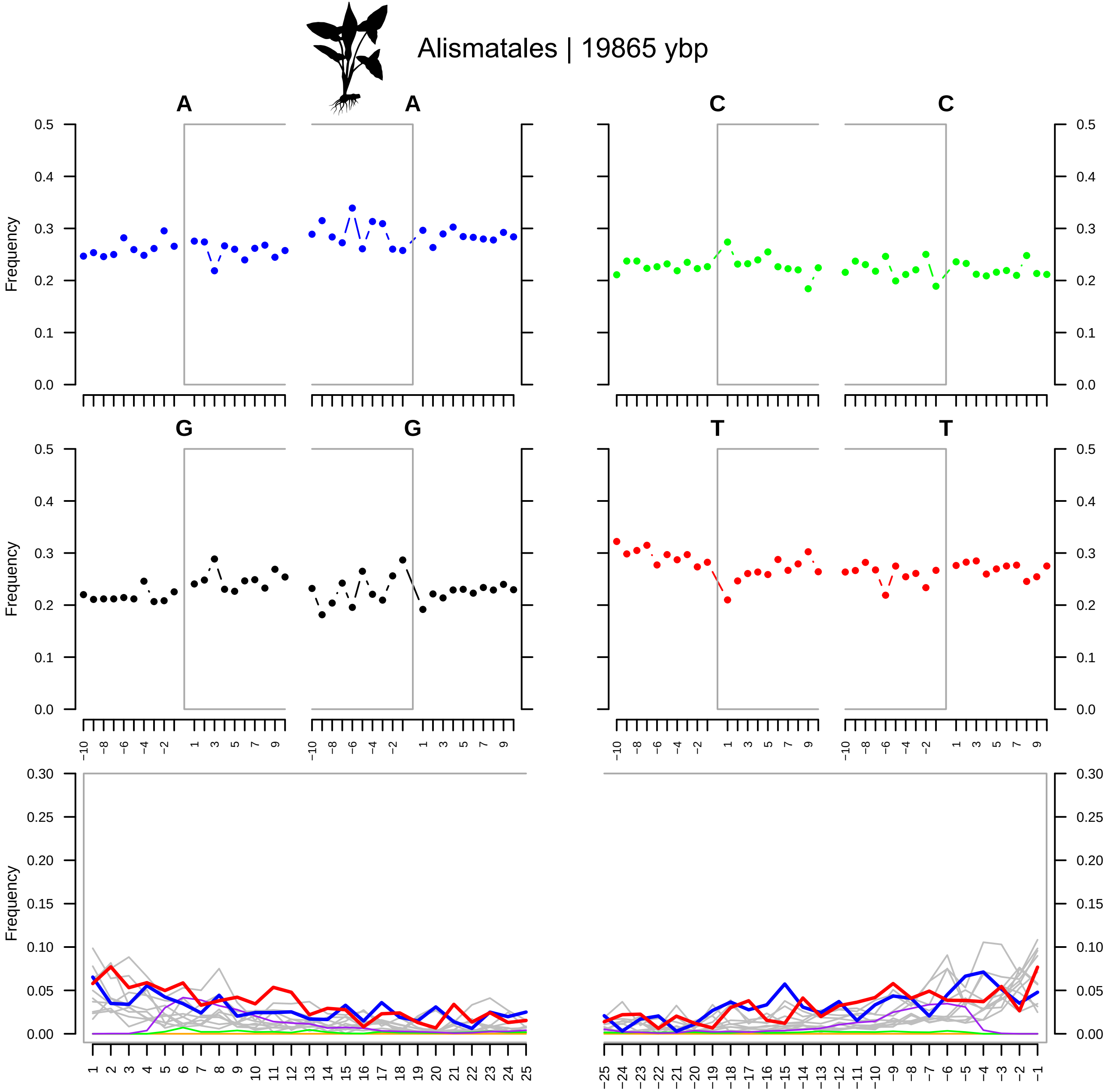

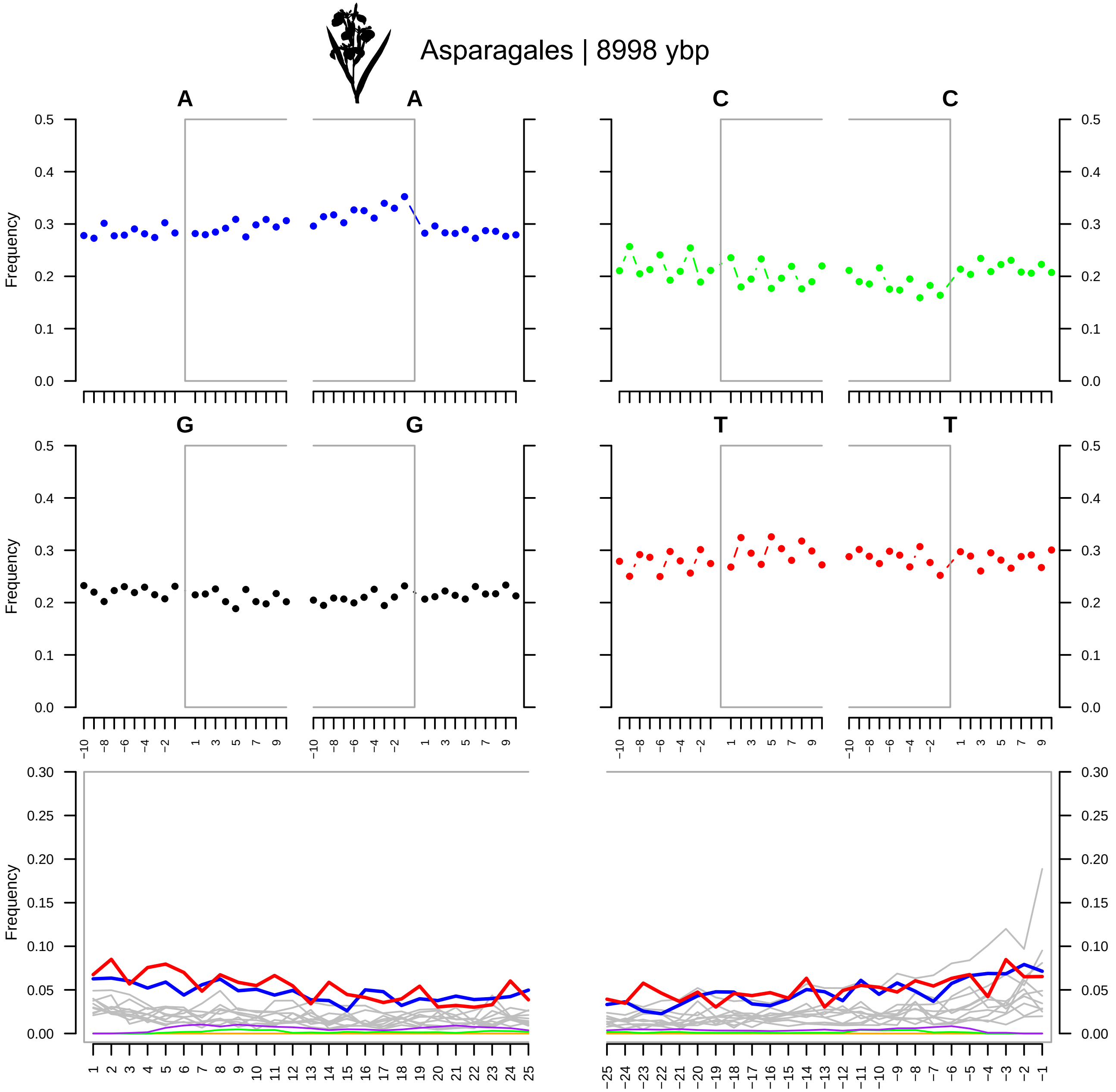
